# 3D genomic capture of regulatory immuno-genetic profiles in COVID-19 patients for prognosis of severe COVID disease outcome

**DOI:** 10.1101/2021.03.14.435295

**Authors:** Ewan Hunter, Christina Koutsothanasi, Adam Wilson, Francisco C. Santos, Matthew Salter, Ryan Powell, Ann Dring, Paulina Brajer, Benedict Egan, Jurjen W. Westra, Aroul Ramadass, William Messer, Amanda Brunton, Zoe Lyski, Rama Vancheeswaran, Andrew Barlow, Dmitri Pchejetski, Peter A. Robbins, Jane Mellor, Alexandre Akoulitchev

## Abstract

Human infection with the SARS-CoV-2 virus leads to coronavirus disease (COVID-19). A striking characteristic of COVID-19 infection in humans is the highly variable host response and the diverse clinical outcomes, ranging from clinically asymptomatic to severe immune reactions leading to hospitalization and death. Here we used a 3D genomic approach to analyse blood samples at the time of COVID diagnosis, from a global cohort of 80 COVID-19 patients, with different degrees of clinical disease outcomes. Using 3D whole genome *EpiSwitch^®^* arrays to generate over 1 million data points per patient, we identified a distinct and measurable set of differences in genomic organization at immune-related loci that demonstrated prognostic power at baseline to stratify patients with mild forms of illness and those with severe forms that required hospitalization and intensive care unit (ICU) support. Further analysis revealed both well established and new COVID-related dysregulated pathways and loci, including innate and adaptive immunity; ACE2; olfactory, Gβγ, Ca^2+^ and nitric oxide (NO) signalling; prostaglandin E2 (PGE2), the acute inflammatory cytokine CCL3, and the T-cell derived chemotactic cytokine CCL5. We identified potential therapeutic agents for mitigation of severe disease outcome, with several already being tested independently, including mTOR inhibitors (rapamycin and tacrolimus) and general immunosuppressants (dexamethasone and hydrocortisone). Machine learning algorithms based on established *EpiSwitch^®^* methodology further identified a subset of 3D genomic changes that could be used as prognostic molecular biomarker leads for the development of a COVID-19 disease severity test.

## Background

As of January 2021, an estimated 85 million people have been infected with the SARS-CoV-2 virus worldwide [1]. Infection with the virus has been shown to induce a wide range of clinical symptoms; some individuals experience asymptomatic or mild disease courses that do not require hospitalization while others develop severe coronavirus disease (COVID-19) triggered by a strong systemic immune response that can lead to acute respiratory failure, thromboembolic phenomena, microvascular disease, viral sepsis and sometimes death [2]. Another early feature associated with more severe disease is significant hypoxemia that commonly occurs in the absence of other systemic symptoms [3]. Several clinical features for disease severity risk following SARS-CoV-2 infection have been identified through epidemiological studies. These include advanced age, male gender, obesity, diabetes, hypertension, the existence of underlying medical conditions such as neurologic disability particularly stroke and renal disease, and immuno-compromised status [4]. However, a subgroup of ‘healthy’ patients without these risk factors nevertheless can develop significant disease leading to increased morbidity and mortality. Identifying the cellular and molecular factors responsible for the heterogeneous disease presentation in COVID-19 is critical for understanding individual disease risk and designing appropriate therapeutic interventions.

Recent studies have identified a central role for the immune system in mediating disease severity following SARS-CoV-2 infection. Individuals with severe disease have higher levels of circulating inflammatory mediators including interleukin (IL)-2, IL-6, IL-7, IL-10, tumour necrosis factor (TNF), granulocyte colony-stimulating factor (G-CSF), monocyte chemoattractant protein-1 (MCP1), and macrophage inflammatory protein 1 alpha (MIP1α) after SARS-CoV-2 infection compared to individuals with mild disease [5, 6]. IL6, in particular, became the focus of attention as a potential disease severity biomarker, as well as a candidate for therapeutic intervention, due to its central role in mediating cytokine storms and the availability of existing targeted therapeutic drugs against it [7, 8]. However, subsequent studies called into question the prominence of IL6 and cytokine storms in SARS-Cov2 infections and several clinical trials of drugs targeting IL6 showed inconsistent results and no clinical benefit, especially for the moderately ill patients [9–13].. However, reports on IL6 treatment with tocilizumab for a specific subgroup of critically ill patients in the first two days of ICU admission demonstrated reduced risk of mortality [14]. While elevated levels of pro-inflammatory cytokines are observed in COVID-19, the exact mechanisms of immune dysfunction leading to severe symptoms remain unclear. As individuals exposed to COVID-19 infections develop a range of acute or long-term systemic inflammatory responses, the ability to identify prognostic profiles for severe clinical cases where patients develop hyperinflammation, do not respond to the standard of care, and progress to needing ICU support is of immediate clinical utility.

The 3D configuration of the genome plays a crucial role in gene regulation [15–17]. The 3D genome acts as the regulatory interface and integration point for multiple inputs: genetic variants and genetic risks, epigenetic modifications, metabolic signals and transcriptional events, influencing cellular phenotype and ultimately clinical outcomes [15, 18, 19]*. EpiSwitch^®^* is a chromosome conformation capture (3C) methodology [20, 21] that has been translated to practice for the discovery of blood-based 3D genomic biomarkers and for patient stratification. To date, 3D genomic biomarkers based on *EpiSwitch^®^* technology have been used to successfully stratify melanoma patients, prognostically stratify patients with fast and slow progressing motor neurone disease, stratify patients with symptomatic and pre-symptomatic Huntington’s disease, diagnose patients with thyroid cancer and various stages of prostate cancer, prognostically stratify patients for treatment in diffuse large B cell lymphoma, and predictively stratify patients with non-small cell lung cancer for response to the PD-L1 immuno-checkpoint inhibitor, avelumab [22–30].

Here we used the *EpiSwitch^®^* Explorer array platform for whole genome profiling of COVID patients at the time of confirmed infection to identify 3D genomic profiles, also known as chromosome conformation signatures (CCS), associated with the development of severe clinical outcomes, requiring ventilation and admission to intensive care units (ICU). After analysing 1.1 million annotated sites across the whole genome for each patient, we found significant and reproducible systemic differences in the 3D genomic profiles of patients presenting different levels of COVID-19 disease severity (mild and severe forms of the disease). To predict disease severity in patients, a subset of prognostic markers at immune-related loci displaying alternative 3D genomic conformations were used to develop a molecular classifier. The markers consistently present at baseline of diagnosis and associated with mild or severe outcomes revealed a regulatory network of biological signalling pathways dysregulated at the level of 3D genome architecture. Analysis of the protein products of the immune-related genomic loci from the predictive marker panel identified a set of existing therapeutic agents that may have clinical utility in managing COVID-19 symptoms. The 3D genomic profile in COVID-19 patients identified through the use of *EpiSwitch^®^* Explorer arrays demonstrated prognostic power and could be further refined using the established *EpiSwitch^®^* methodology to develop a PCR-based test for prognosis of severe outcomes in COVID-19 [22–30].

## Materials and Methods

### Patient characteristics

Clinical PBMC and whole blood samples from consented patients were obtained from academic collaborators and commercial sources. A total of 80 patients from 4 sample cohorts were used in this study, comprising a multinational set of COVID-19 cases: asymptomatic, mild hospitalized and severe (ICU support) - from the United Kingdom, the United States, and Peru. All samples were collected at the time of diagnosis of COVID infection with PCR test. Patients were then observed over the period of up to several weeks for clinical manifestations of COVID disease. The age of the patients ranges from 24 to 95, with median at 70 years (**Table 1**).

**Table 1:** Patient information used in all the cohorts.

| Cohort | Gender | Age | Race | Ethnicity | Country Collected | COVID Severity |
| --- | --- | --- | --- | --- | --- | --- |
| 1 | Male | 80 | Other | Any other ethnic group | United Kingdom | Mild |
| 1 | Female | 85 | White | White/Any Other White Background | United Kingdom | Mild |
| 1 | Male | 75 | White | White British | United Kingdom | Mild |
| 1 | Male | 36 | Asian | Asian / Any Other Asian Background | United Kingdom | Severe |
| 1 | Male | 70 | White | White British | United Kingdom | Severe |
| 1 | Male | 87 | White | White - British | United Kingdom | Mild |
| 1 | Female | 89 | White | White/Any Other White Background | United Kingdom | Mild |
| 1 | Female | 95 | White | White/Any Other White Background | United Kingdom | Mild |
| 1 | Male | 49 | White | White/Any Other White Background | United Kingdom | Severe |
| 1 | Female | 83 | Other | Any other ethnic group | United Kingdom | Severe |
| 1 | Male | 73 | White | White/Any Other White Background | United Kingdom | Severe |
| 2 | Male | 62 | White | Non-Hispanic | United States | Asymptomatic |
| 2 | Male | 55 | White | Non-Hispanic | United States | Mild/Moderate |
| 2 | Female | 62 | Black | Non-Hispanic | United States | Asymptomatic |
| 2 | Male | 63 | White | Non-Hispanic | United States | Mild/Moderate |
| 2 | Male | 71 | White | Non-Hispanic | United States | Mild/Moderate |
| 2 | Female | 24 | Black | Non-Hispanic | United States | Mild/Moderate |
| 2 | Female | 67 | Black | Non-Hispanic | United States | Mild/Moderate |
| 2 | Male | 65 | White | Non-Hispanic | United States | Mild/Moderate |
| 2 | Female | 57 | Black | Non-Hispanic | United States | Mild/Moderate |
| 2 | Female | 30 | Black | Unknown | United States | Mild/Moderate |
| 2 | Male | 71 | White | Non-Hispanic | United States | Asymptomatic |
| 2 | Male | 59 | White | Non-Hispanic | United States | Mild/Moderate |
| 3 | Male | 67 | White | Non-Hispanic | United States | Severe |
| 3 | Female | 62 | White | Non-Hispanic | United States | Mild |
| 3 | Male | 87 | White | Non-Hispanic | United States | Mild |
| 3 | Female | 82 | White | Non-Hispanic | United States | Severe |
| 3 | Female | 47 | White | Hispanic | United States | Severe |
| 3 | Female | 47 | White | Non-Hispanic | United States | Severe |
| 3 | Female | 27 | White | Non-Hispanic | United States | Asymptomatic |
| 3 | Female | Unk<sup>a</sup> | White | Non-Hispanic | United States | Asymptomatic |
| 3 | Female | 52 | White | Non-Hispanic | United States | Asymptomatic |
| 3 | Female | 42 | White | Non-Hispanic | United States | Mild - Not hospitalised |
| 3 | Male | 72 | White | Non-Hispanic | United States | Mild - Not hospitalised |
| 3 | Male | 67 | White | Non-Hispanic | United States | Mild |
| 3 | Female | Unk | Black | Non-Hispanic | United States | Severe |
| 3 | Female | Unk | Unk | Unk | United States | Mild - Not hospitalised |
| 3 | Female | 38 | White | Non-Hispanic | United States | Asymptomatic |
| 4 | Male | 84 | N/A | Hispanic | Peru | ICU |
| 4 | Male | 56 | N/A | Hispanic | Peru | Hospitalised |
| 4 | Female | 82 | N/A | Hispanic | Peru | Hospitalised |
| 4 | Male | 88 | N/A | Hispanic | Peru | Hospitalised |
| 4 | Male | 78 | N/A | Hispanic | Peru | Hospitalised |
| 4 | Male | 79 | N/A | Hispanic | Peru | Hospitalised |
| 4 | Male | 70 | N/A | Hispanic | Peru | ICU |
| 4 | Male | 64 | N/A | Hispanic | Peru | ICU |
| 4 | Male | 72 | N/A | Hispanic | Peru | ICU |
| 4 | Male | 70 | N/A | Hispanic | Peru | Hospitalised |
| 4 | Female | 67 | N/A | Hispanic | Peru | ICU |
| 4 | Female | 57 | N/A | Hispanic | Peru | Hospitalised |
| 4 | Female | 84 | N/A | Hispanic | Peru | ICU |
| 4 | Male | 49 | N/A | Hispanic | Peru | Hospitalised |
| 4 | Female | 83 | N/A | Hispanic | Peru | ICU |
| 4 | Male | 88 | N/A | Hispanic | Peru | ICU |
| 4 | Female | 66 | N/A | Hispanic | Peru | ICU |
| 4 | Female | 77 | N/A | Hispanic | Peru | ICU |
| 4 | Male | 89 | N/A | Hispanic | Peru | ICU |
| 4 | Male | 70 | N/A | Hispanic | Peru | Hospitalised |
| 4 | Male | 78 | N/A | Hispanic | Peru | ICU |
| 4 | Female | 77 | N/A | Hispanic | Peru | Hospitalised |
| 4 | Female | 83 | N/A | Hispanic | Peru | ICU |
| 4 | Male | 69 | N/A | Hispanic | Peru | Hospitalised |
| 4 | Female | 63 | N/A | Hispanic | Peru | ICU |
| 4 | Female | 82 | N/A | Hispanic | Peru | ICU |
| 4 | Male | 60 | N/A | Hispanic | Peru | Hospitalised |
| 4 | Male | 78 | N/A | Hispanic | Peru | Hospitalised |
| 4 | Male | 75 | N/A | Hispanic | Peru | ICU |
| 4 | Male | 92 | N/A | Hispanic | Peru | ICU |
| 4 | Female | 64 | N/A | Hispanic | Peru | Hospitalised |
| 4 | Male | 75 | N/A | Hispanic | Peru | Hospitalised |
| 4 | Male | 64 | N/A | Hispanic | Peru | ICU |
| 4 | Female | 79 | N/A | Hispanic | Peru | ICU |
| 4 | Male | 78 | N/A | Hispanic | Peru | Hospitalised |
| 4 | Female | 77 | N/A | Hispanic | Peru | ICU |
| 4 | Male | 86 | N/A | Hispanic | Peru | Hospitalised |
| 4 | Female | 77 | N/A | Hispanic | Peru | ICU |
| 4 | Male | 68 | N/A | Hispanic | Peru | ICU |
| 4 | Male | 63 | N/A | Hispanic | Peru | Hospitalised |
| 4 | Male | 68 | N/A | Hispanic | Peru | ICU |
| 4 | Male | 71 | N/A | Hispanic | Peru | ICU |

<sup>a</sup> *Unk: Unknown.*

### Array design

Custom microarrays were designed using the *EpiSwitch^®^* pattern recognition algorithm, which operates on Bayesian-modelling and provides a probabilistic score that a region is involved in long-range chromatin interactions. It was used to annotate the GRCh38 human genome assembly across ∼1.1 million sites with the potential to form long-range chromosome conformations [22–30]. The most probable interactions were identified and filtered on probabilistic score and proximity to protein, long non-coding RNA, or microRNA coding sequences. Predicted interactions were limited to *EpiSwitch^®^* sites greater than 10 kb and less than 300 kb apart. Repeat masking and sequence analysis was used to ensure unique marker sequences for each interaction. The *EpiSwitch^®^* Explorer array (Agilent Technologies, Product Code X-HS-AC-02), containing 60-mer oligonucleotide probes was designed to interrogate potential 3D genomic interactions. In total, 964,631 experimental probes and 2,500 control probes were added to a 1 x 1 M CGH microarray slide design. The experimental probes were placed on the design in singlicate with the controls in groups of 250. The control probes consisted of six different *EpiSwitch^®^* interactions that are generated during the extraction processes and used for monitoring library quality. A further four external inline control probe designs were added to detect non-human (*Arabidopsis thaliana*) spike in DNA added during the sample labelling protocol to provide a standard curve and control for labelling. The external spike DNA consists of 400 bp ssDNA fragments from genomic regions of *A. thaliana*. Array-based comparisons were performed described previously, with the modification of only one sample being hybridised to each array slide in the Cy3 channel [22–30].

### Preparation of genomic templates

Chromatin with intact chromosome conformations from each blood sample was extracted using the *EpiSwitch^®^* Explorer Array Kit following the manufacturer’s instructions (Oxford BioDynamics Plc) [22–30]. The *EpiSwitch^®^* Explorer arrays were performed as published previously, with the modification of only one sample being hybridised to each array slide in the Cy3 channel. *EpiSwitch^®^* Explorer arrays, based on Agilent SureSelect array platform, allow for the highly reproducible, non-biased interrogation of ∼1.1 million anchor sites for 3D genomic interactions (964,631 experimental probes and 2500 control probes).

### Statistical analysis

The four COVID cohorts were normalised by background correction and quantile normalisation, using the *EpiSwitch^®^* R analytic package, which is built on the Limma and dplyr libraries. The four datasets were then combined into one sample set containing 80 samples. Data was corrected for batch effects using ComBat R script. Parametric (Limma R library, Linear Regression) and non-parametric (*EpiSwitch^®^* RankProd R library) statistical methods were performed to identify 3D genomic changes that demonstrated a difference in abundance between the Mild and Severe COVID-19 classes. Asymptomatic patients (10 samples) were excluded from this analysis. The resulting data from both procedures were further filtered based on adjusted p-value (FDR correction) and abundance scores (AS). Only 3D genomic markers with adjusted p-value <=0.05 and AS −1.1<= or >=1.1 were selected. Both filtered lists from Limma and RankProd analysis were compared and the intersection of the two lists was selected for further processing.

### Genome mapping and linear discriminant analysis

1000 3D genomic markers from the statistically filtered list with the greatest and lowest abundance scores were selected for genome mapping. Mapping was carried out using Bedtools *closest* function for the 3 closest protein coding loci (Gencode v33). The resulting list of ‘Severe’ and ‘Mild’ 3D genomic markers were further annotated for relatedness to immunological processes using the ‘immune process’ annotation from Gene Ontology and gene lists for immune aging and trained immunity [31–33]. Significant 3D genomic markers with associated protein coding loci involved in immune processes were then ordered by adjusted p-value (adj.P.Val), then abundance score. The top 100 3D genomic markers from this combined filter were then utilized for linear discriminant analysis (LDA) using the MASS library and visualized using the ggplot2 package in R.

### Machine learning

Two approaches were selected for model building, linear discriminant analysis (LDA) and eXtreme Gradient Boosting (XGBoost). Both procedures were performed using R, with caret, XGBoost, and SHAPforxgboost libraries for the XGBoost model and MASS library for LDA model.

### Biological network and drug target analysis

Network analysis for functional/biological relevance of the 3D genomic markers was performed using the Hallmark Gene Sets and BioCarta and Reactome Canonical Pathway gene sets from the Molecular Signatures Database (MSigDB) [34]. Protein interaction networks were generated using the Search Tool for the Retrieval of Interacting proteins (STRING) database [35]. Candidate drugs were identified using the GeneAnalytics platform (geneanalytics.genecards.org) [36].

## Results

### Array-based profiling of COVID-19 patient cohorts

We used whole-genome *EpiSwitch^®^* Explorer arrays to screen peripheral blood mononuclear cells (PBMC) samples collected at the time of confirmed COVID-19 infection from 38 patients in three independent cohorts from the US and the UK. Interestingly, all three cohorts showed separation by principal component analysis (PCA) for Mild or Severe (requiring ICU admission) outcomes without pre-selection or reduction of the array markers (**Figure 1**), suggesting that 3D genomic profiles associated with different clinical outcomes existed and could be distinguished.

**Figure 1.**
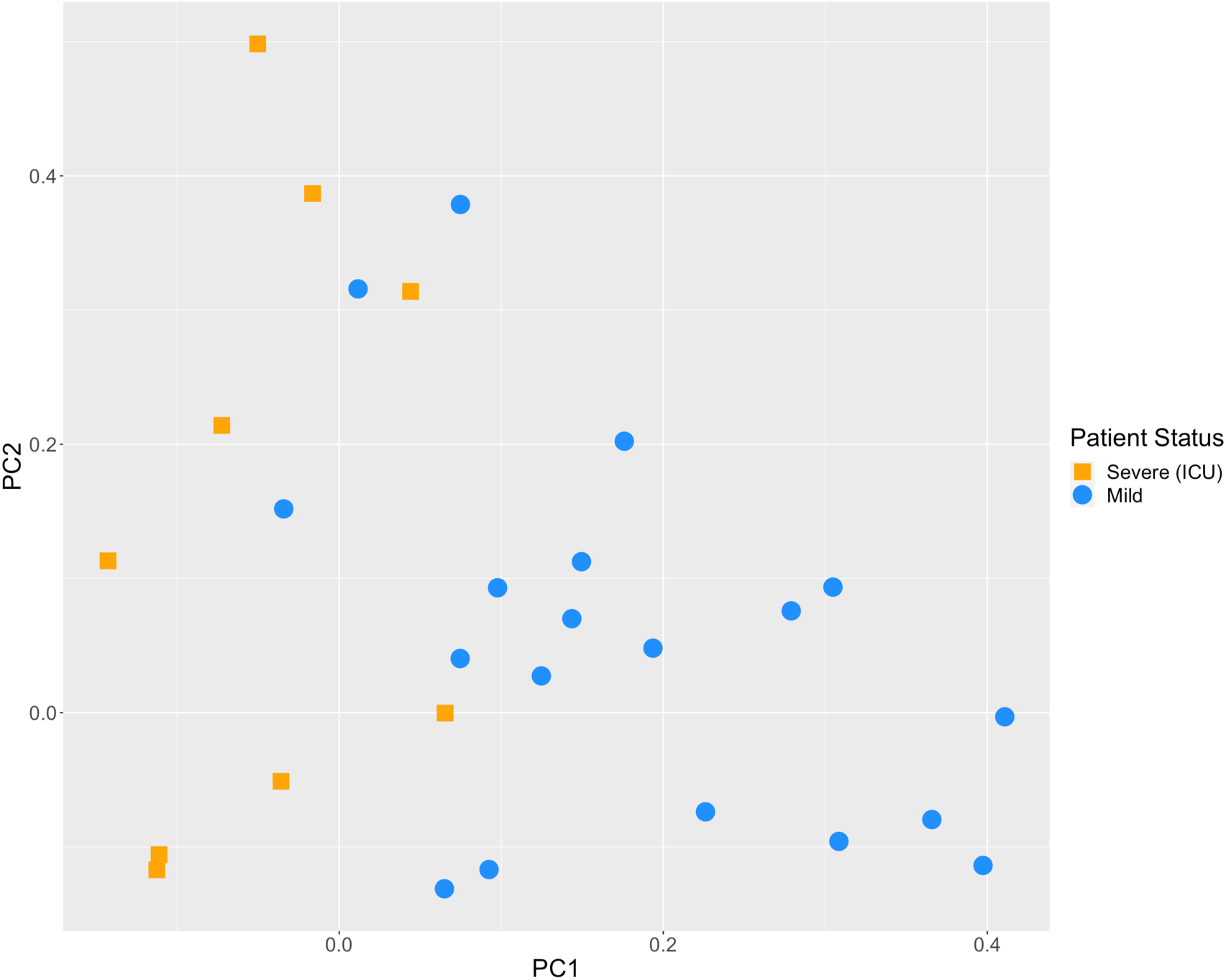
PCA plot of three independent COVID-19 cohorts from the UK and USA for Mild and Severe (ICU) disease outcomes. The PCA plot of COVID-19 patients that exhibited mild disease outcomes (blue circles) and severe disease outcomes requiring ICU admission (orange squares) is based on whole genome profiling of all 964 thousand CCS markers screened, without any marker reduction.

### Identification of the top prognostic 3D genomic markers for Severe & Mild COVID-19 disease outcomes

To evaluate the biological relevance of the observed separation of Mild and Severe COVID-19 outcomes by 3D genomic markers, we focused on the top 750 markers prognostically associated with each of the two outcomes. The statistical testing employed in this study to determine statistically significant 3D genomic markers benefits from using both parametric testing (Limma) and non-parametric testing (*EpiSwitch^®^* RankProd), both procedures that correct for multiple testing by using False Discovery Rate (FDR) corrections. The RankProd approach also has a resampling step to control for random rank importance, adding another layer of statistical stringency in marker selection when testing a large number of possibilities. The top markers selected were filtered based on an adjusted (FDR) P value <=0.05, and high abundance scores (AS), −1.1<= or >=1.1. These criteria were applied to the parametric and non-parametric statistical sets, producing the top 750 intersecting markers. Similar approaches and thresholds for FDR cut-offs were utilized in previously published biomarker development studies using *EpiSwitch^®^* [22–30].

We next evaluated the top 750 3D genomic markers, identified from 964,631 whole genome screened cis-interactions, according to the functional roles of the genes encoded across their genomic locations using Hallmark Gene Sets as well as BioCarta and Reactome canonical pathway analysis (**Figure 2**). The list of affected pathways and corresponding genetic loci with individual 3D genomic changes is provided in **Supplementary Tables 1a, b**. When evaluating the biological function of the genes within the genomic regions identified as being dysregulated between patients who developed mild or severe outcomes in COVID-19, a number of biological pathways with known associations to COVID-19 were identified including; the olfactory signalling pathway ACE2, innate and adaptive immune systems, IL6 and JAK-STAT signalling, calcium signalling, (NO) nitric oxide signalling, coagulation, complement, interferon gamma response, TGF beta signalling, TNF alpha signalling, and apoptosis [37–42]. In addition, the MSP-RON systemic inflammatory response was identified as a potentially novel signalling pathway dysregulated in COVID-19.

**Figure 2.**
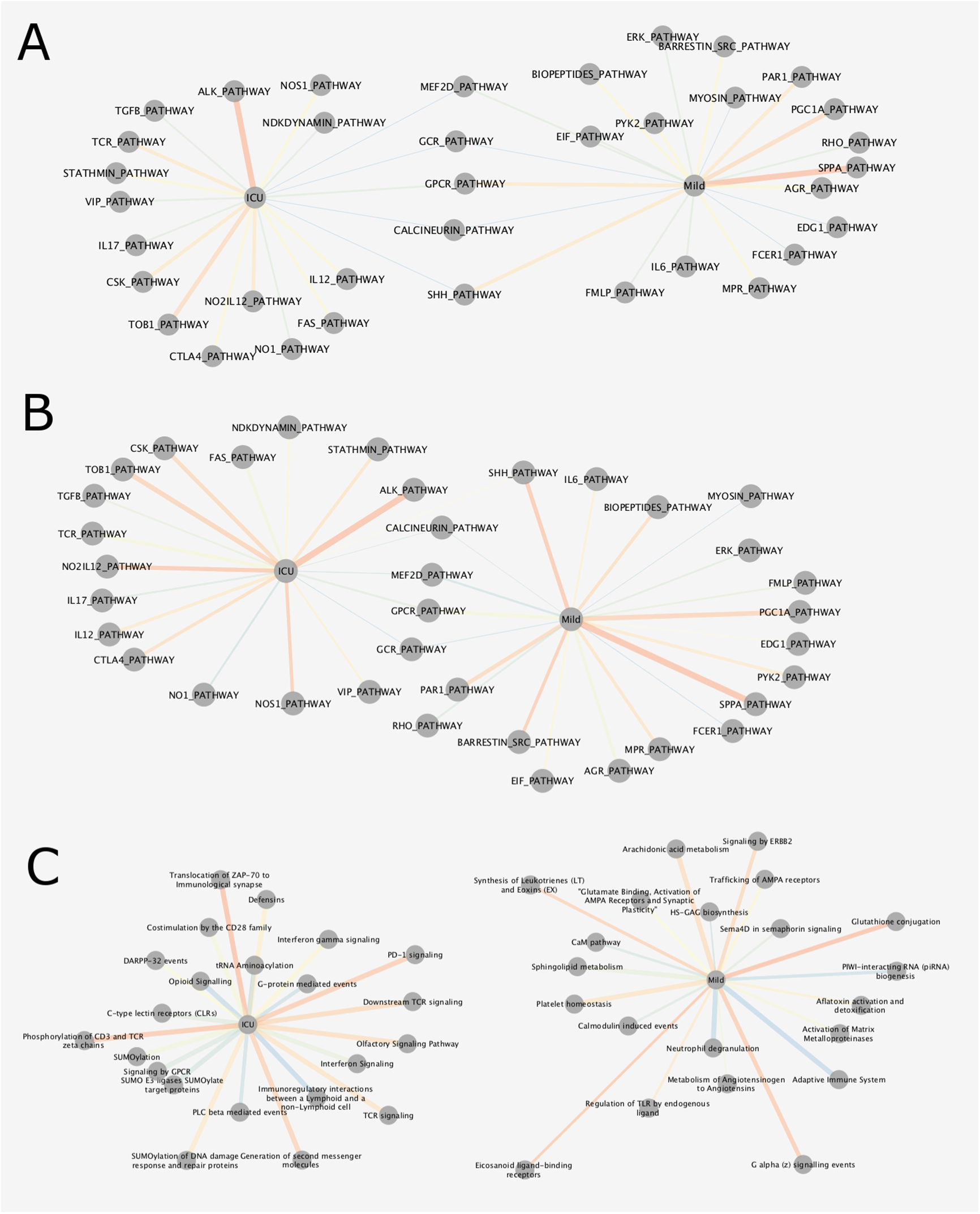
Mapping of top 3D genomic markers to biological pathways. Gene set analysis of the top 750 3D genomic markers separating Mild and Severe (ICU) COVID-19 outcomes using Hallmark (A), BioCarta (B), and Reactome (C) gene and canonical pathway lists. Thickness of edges indicates the number of mapping dysregulated genes and color indicates statistical rank (orange - high rank; blue - lower rank). See also **Supplementary Tables 1a, b.**

Interestingly, SARS-CoV-2 has been reported to drive hyperactivation of CD4+ T-cells and immune paralysis (due to loss of FOXP3 negative feedback) to promote pathogenesis of disease [43]. CD25+ (IL2 alpha receptor) FOXP3- are considered hyperactive T-cells and fail to differentiate into regulatory T-cells (Tregs) and produce Furin to promote viral entry into lung epithelial cells [43]. CD28 (see Fig.2C, Reactome, ICU) may function with IL2 to mediate the feedbacks necessary to repress a potentially overstimulated immune response in COVID19. With CD25 being the IL2 alpha receptor, IL-2 acts as a potent growth factors for CD25-expressing activated T cells (see Fig.2A Hallmark, ICU). The prevalence of both IL2 and CD25 indicates that a positive feedback loop for T-cell activation is established in severe COVID-19 leading to the production of multiple effector cytokines. This may be because of a reduction of FOXP3-mediated negative regulation to allow functional Tregs to be produced. The CD25+ T-cells in severe patients are likely to die partly by cytokine deprivation or become hyperactivated in severe disease – i.e. FOXP3 negative cells may become ex-Tregs or hyperactivated T-cells (leading to T cell paralysis). These abnormally activated T-cells produce Furin which activates the Spike pro-protein cleavage and promotes viral entry into cells.

Regarding the immune checkpoints, IL2 expression activates FOXP3 and prolonged activation results in the expression of immune checkpoints such as CTLA-4 and FOXP3, which represses transcription of effector cytokines, suppression of T-cell responses and resolution of inflammation (i.e. in normal cells). In severe COVID-19, hyperactivated macrophages may present antigens to CD4+ T-cells which are activated and differentiate into CD25+, IL10R+ early activated T-cells which produced IL10 rather than IL2 and there is no Foxp3 expression to start the negative feedback. IL10 further enhances the activation of CD25+ T-cells which express immune checkpoints, multiple cytokines and Furin. Multifaceted Th differentiation leads to unfocused T-cell responses and paralyses the T-cell system.

The nucleocapsid (N) protein of the SARS-Cov-1 virus is sumoylated (see Fig.2C Reactome, ICU) and binds to hUbc9, a ubiquitin conjugating enzyme of the sumoylation system [44]. SARS-CoV-2 N protein is likely to be sumoylated at several sites included K62. This pathway is a potential target for treatments as SUMOylation is required for homo-oligomerisation and self-association of the N protein required for the formation of viral RNP and nucleocapsid assembly.

PD-L1 expression in severe COVID-19 patients (see Fig.2C Reactome, ICU) is likely to be linked to immunosuppressive phenotypes in innate immune cells and to support lymphopenia through apoptosis of lymphocytes [45]. It is possible that PD-1 signalling is not able to control hyperactivated T cells and resolution of hyperinflammatory stage. It remains to be investigated if PD-L1 expression on lung epithelia may also regulate PD-1 expressing T-cells (as shown for influenza and Rous sarcoma viruses [46, 47].

Oxidative phosphorylation (see Fig. 2A Hallmark, ICU) may prove to be the link between the metabolic state of cells in people with predisposing conditions (T1D, heart attacks, obesity, use of steroids, etc), and dysregulation of the homeostasis of CD25+ T-cells and Fox3p expressing Tregs. Activation of Tregs is impaired in Type1 diabetics, but is also reduced in severe COVID patients [48, 49]. FOXP3 expression is reduced in CD25+ CD4+ T-cells in patients who have had heart attacks [50]. Leptin released from adipocytes also prevents CD25+CD4+ T-cell proliferation [51]. T-cell activation is dependent on glycolysis and oxidative phosphorylation while Treg differentiation is more dependent on oxidative phosphorylation and inhibited by glycolysis [52, 53]. This could be because of the hypoxic lung in severe COVID which leads to higher levels of glycolysis – hence reduced Treg differentiation – via HIF-1alpha activation (which mediates glycolysis and so promotes degradation of FoxP3 proteins (reduced feedback loop blocking Treg differentiation). Type 1 interferons (see Fig.2A Hallmark, Mild) and downstream pathways are suppressed in severe patients (i.e. lower levels of IFT1,2,3 and IF1TM1), with lower levels of TNF ligands TRAIL, LIGHT and surface proteins SLAMF1, KLRB1, all of which have roles in viral infections [43, 49].

The profound hypoxia associated with more severe disease may well result from viral damage to hypoxic pulmonary vasoconstriction, which is a protective mechanism that diverts blood flow towards the healthier regions of the lung where oxygen uptake can still occur [54]. The regulation of blood flow within the lung is dependent on both Ca^2+^ signalling and NO. The mechanisms associated with acute hypoxia signalling are not understood, but an interesting link has been made in the carotid body between this mechanism and the olfactory receptor Olfr78 [55]. In fact, *EpiSwitch* array analysis identified 3 statistically significant CCS markers at the Olfr78/OR51E2 locus. A number of other features relating to hypoxia in SARS-CoV-2 infection may be caused by viral infection of carotid body type 1 cells [56].

Olfactory signalling (Reactome ICU) and NO (Biocarta ICU) pathways are linked, as NO acts as a neurotransmitter involved in neural olfactory processes in the central nervous system. and also inhibits viral replication [57, 58] With the loss of smell both sustentacular cells and basal cells appear to be affected, and both express ACE2 (the receptor for the SARS-CoV-2 virus) and TMPRSS2 (a serine protease controlling viral entry into cells) [59].

### Linear Discriminant Analysis across COVID-19 patient cohorts

The unbiased whole genome array screening on three independent cohorts of COVID patients, coupled with the pathway analyses on the top 750 markers, strongly supported immune related genomic loci and pathways associated with different clinical outcomes of COVID-19. To refine this further, we added a fourth blood cohort of hospitalized COVID-19 patients from Lima, Peru which at the time of collection had one of the highest COVID-19 fatality rates in the world (3.5%). Of the 42 hospitalized patients in this cohort, 18 remained on the ward with mild disease and 26 progressed to ICU support. Thus, when combined with the 38 patients in the first 3 cohorts, a total of 80 patients who were screened by the whole genome array were used, providing 77.3 million data points from patients clinically assessed as Asymptomatic (7), Mild (40) and Severe (35). With the focus on prognosis of severe (ICU) outcomes, we reduced our analysis to the top 100 immuno-genetic components of the 3D genomic markers (see **Materials and Methods**) statistically associated with Severe (ICU) outcome in clinical annotations (**Table 2**). Next, this data was subject to Linear Discriminant Analysis (LDA). By LDA, the top 100 Severe (ICU) markers were able to demonstrate statistically significant difference for patients with different clinical outcomes - asymptomatic, mild and severe (ICU) (**Figure 3**).

**Figure 3.**
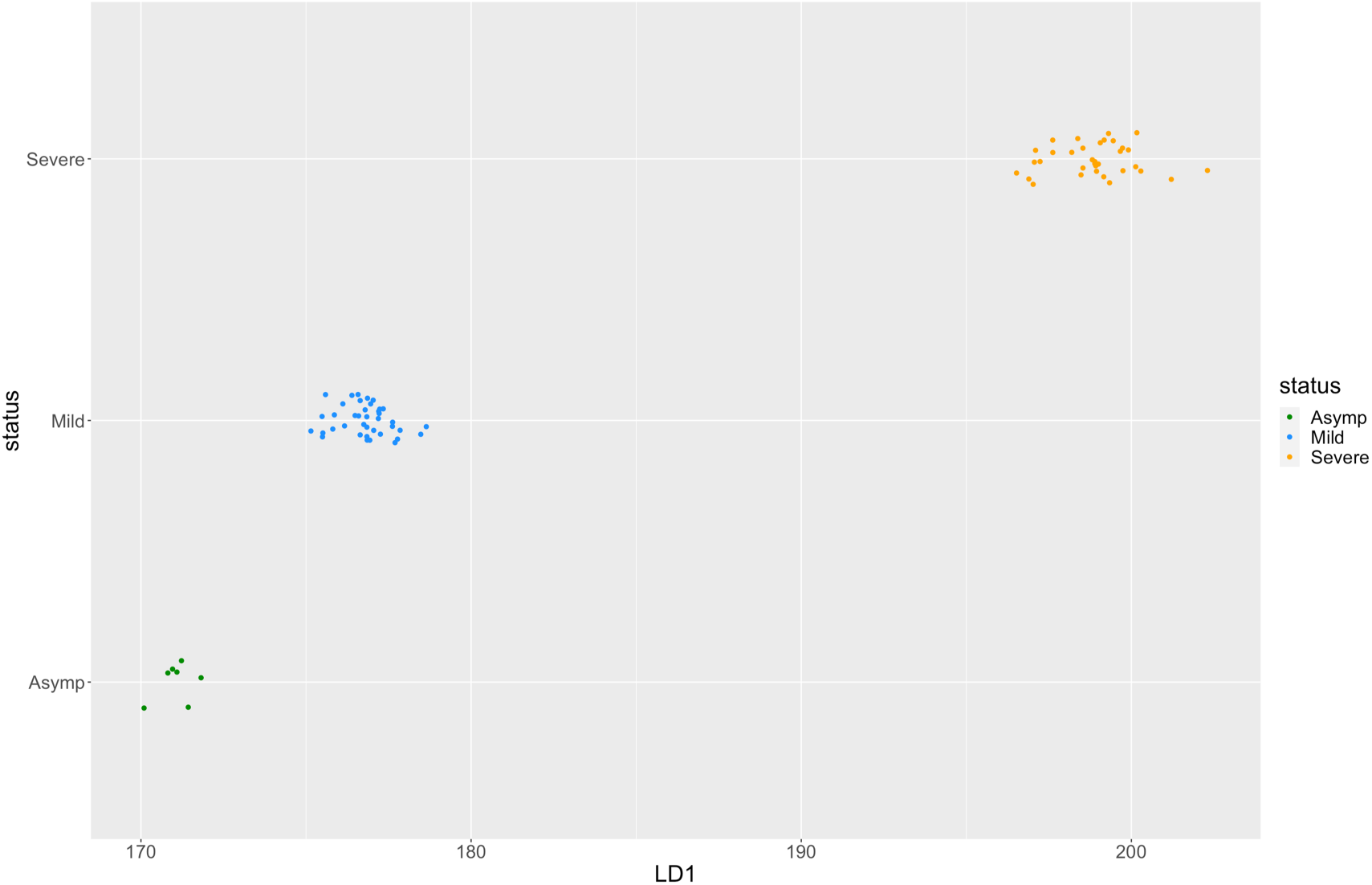
Linear Discriminant Analysis (LDA) for COVID-19 disease severity. The top 100 immune-related 3D genomic markers associated with Severe (ICU) clinical outcomes were used to characterize 80 COVID-19 patients with different disease severity levels ranging from asymptomatic (Asymptomatic, blue circles), Mild presentations of disease (green circles) and this with Severe presentations (red circles). See also **Table 2 a,b.** (Y axis - patient categories; X axis - Linear Discriminant Coordinate 1 (LD1).

**Table 2a:**
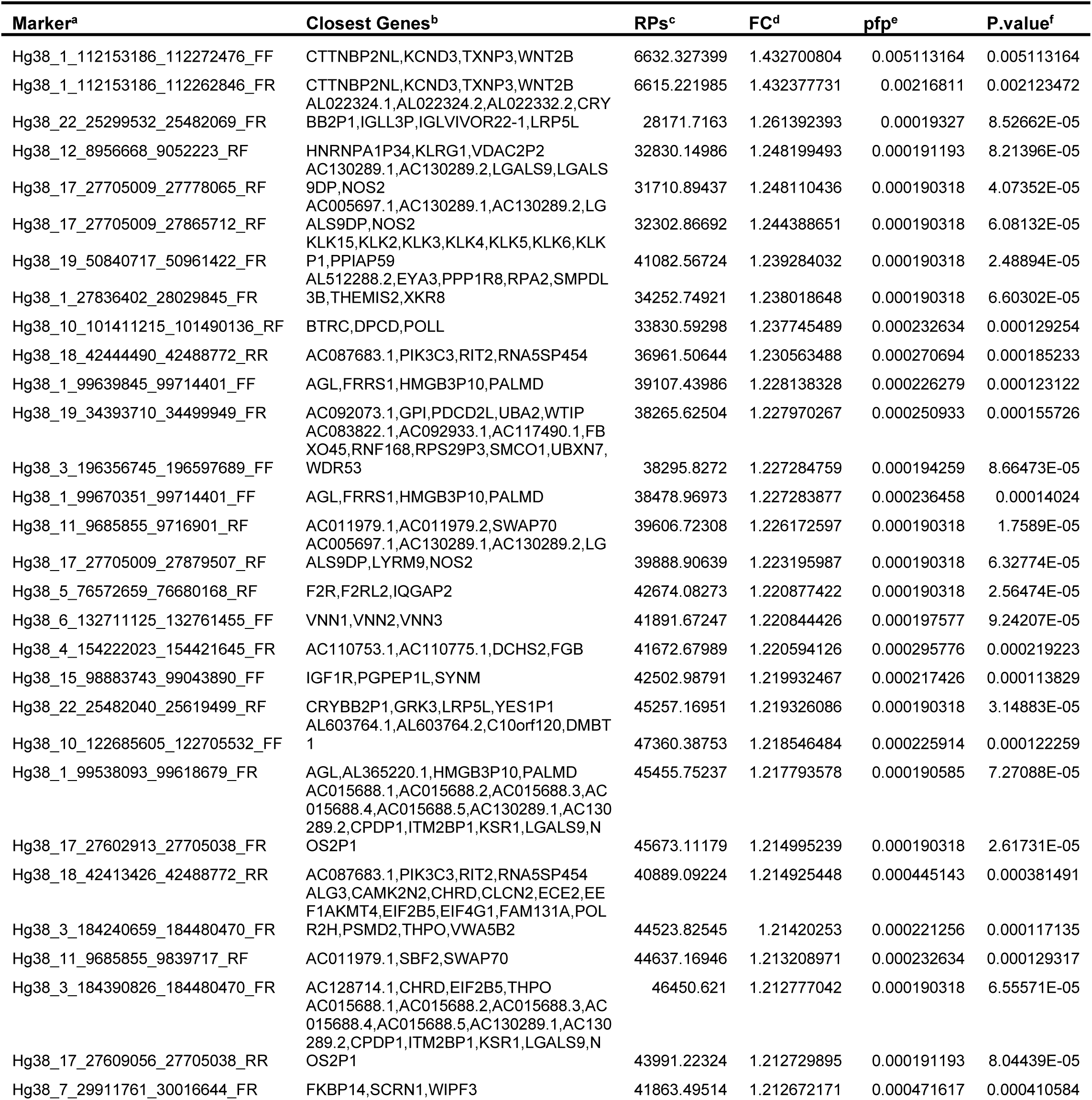
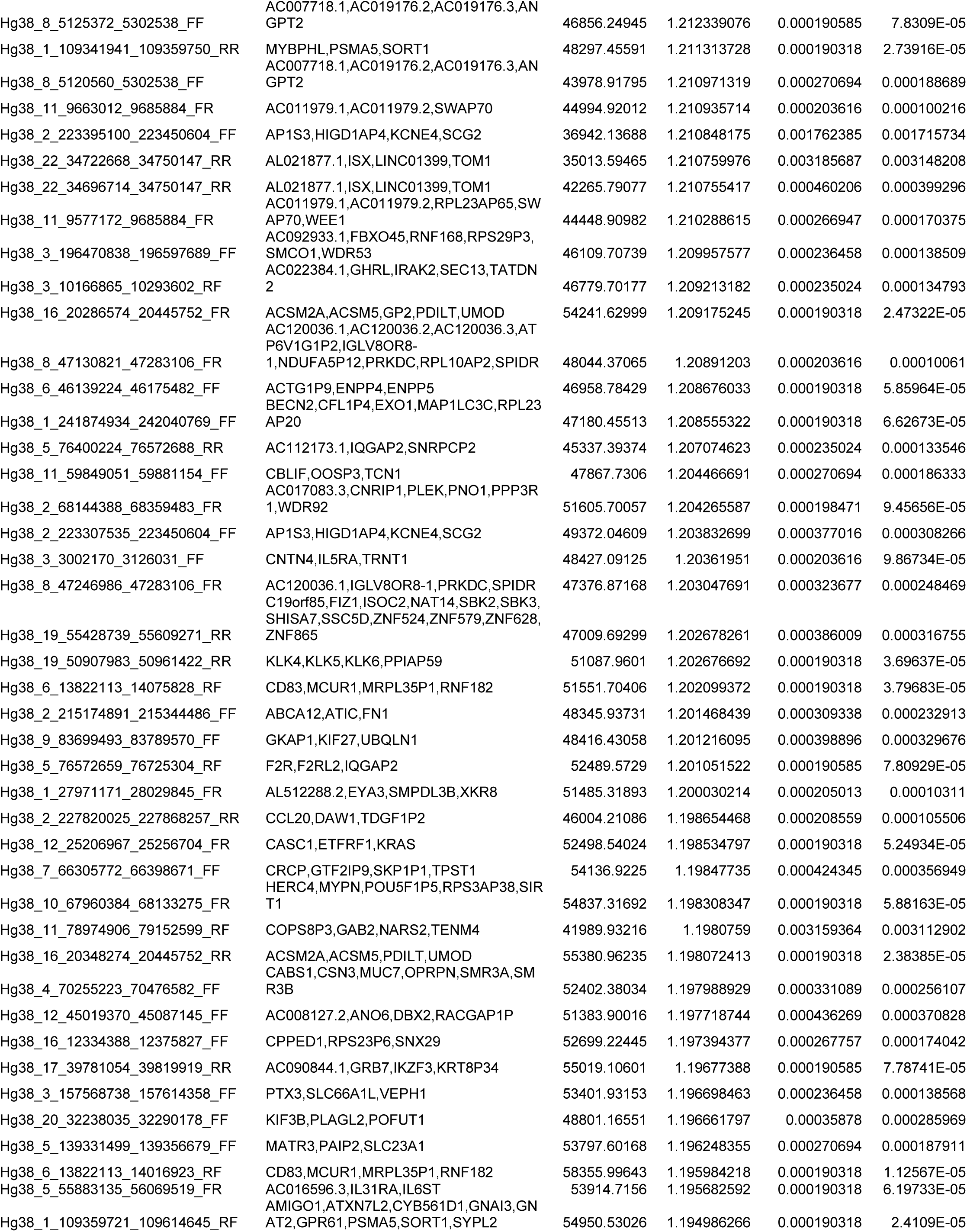
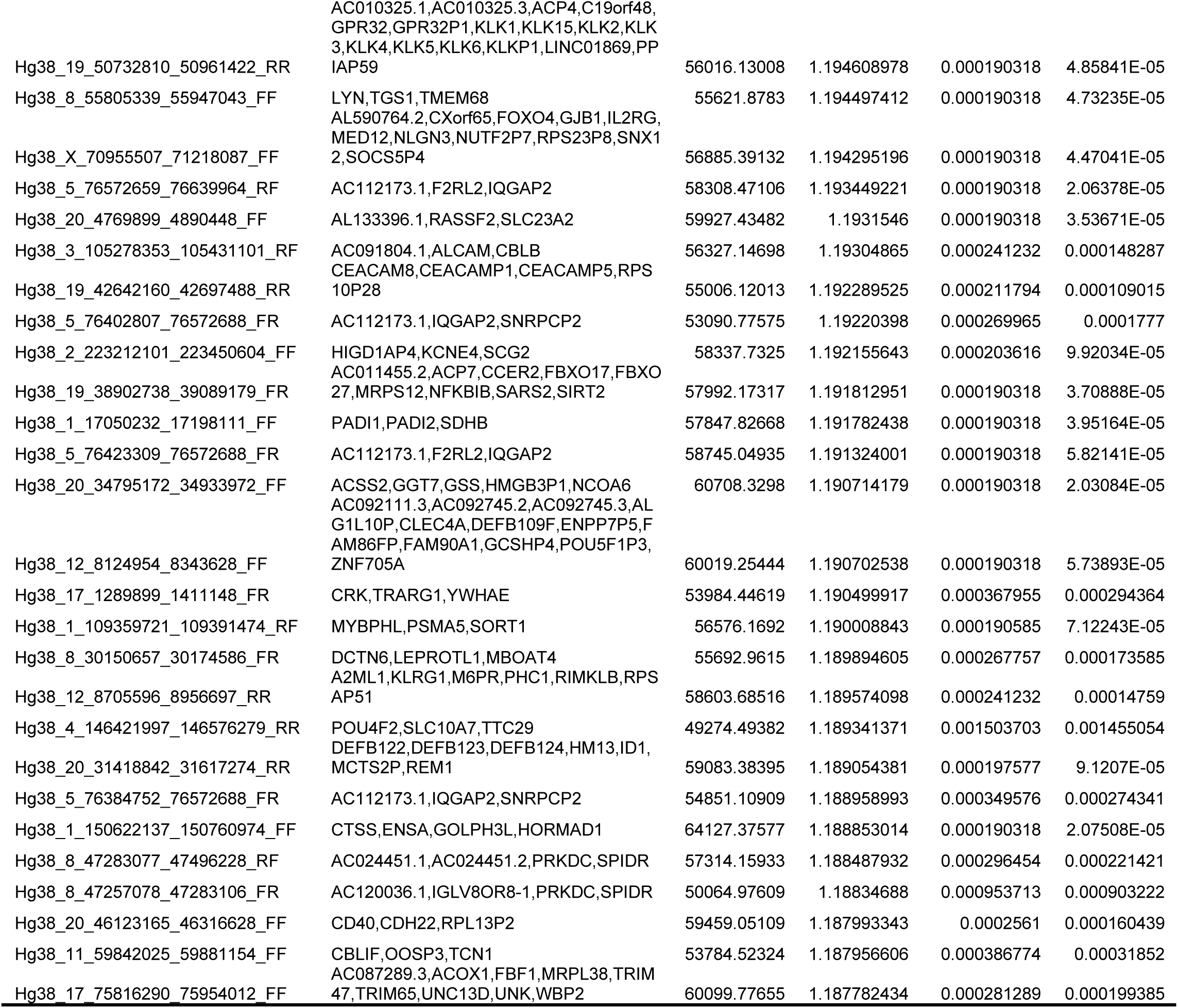
Top 100 3D genomic markers associated with Severe (ICU) clinical outcomes.

**Table 2b:**
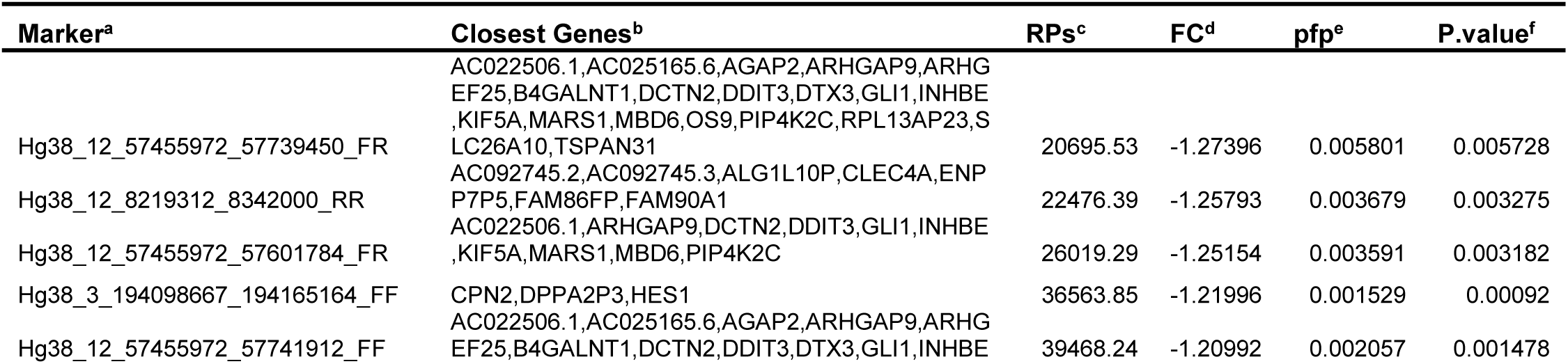
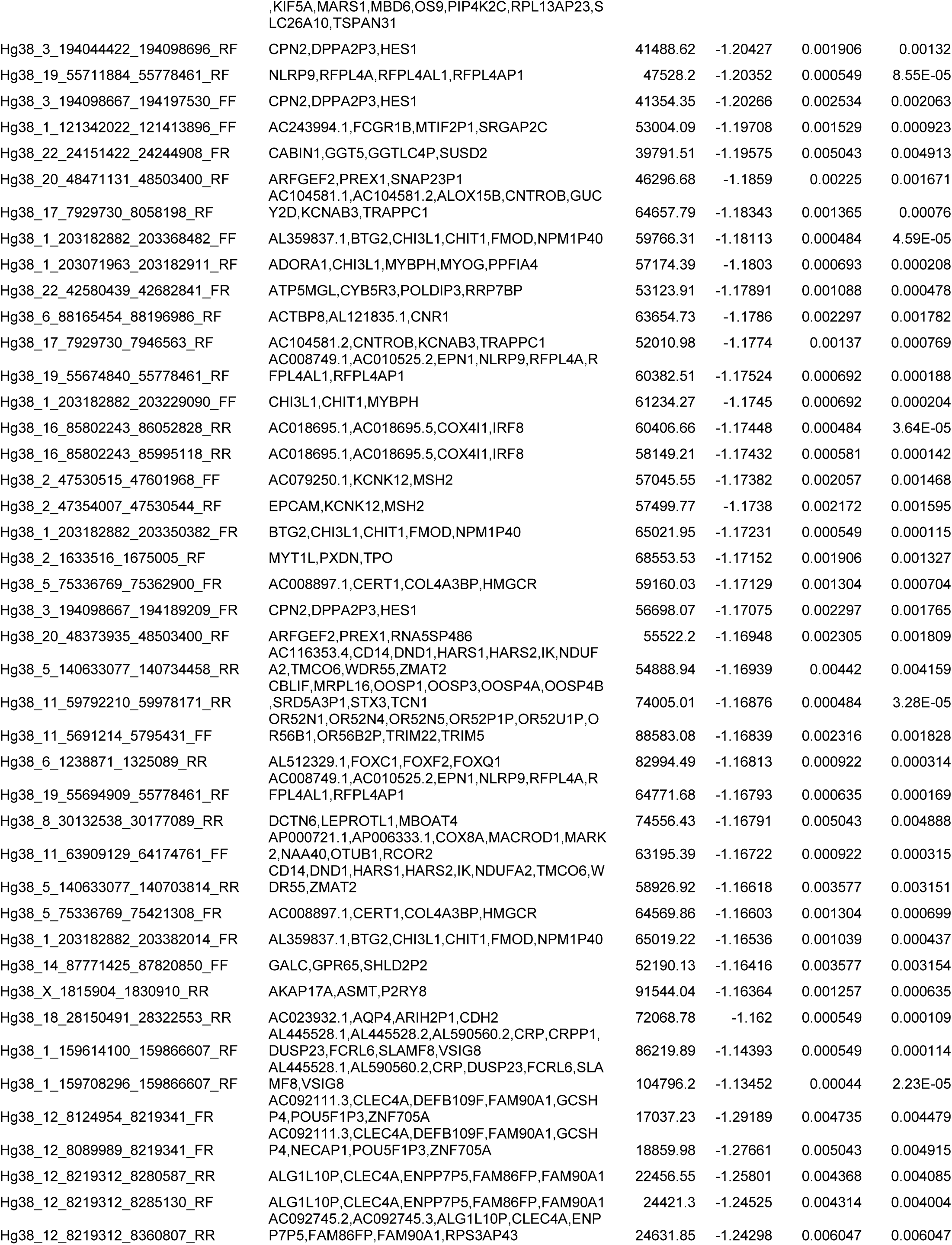
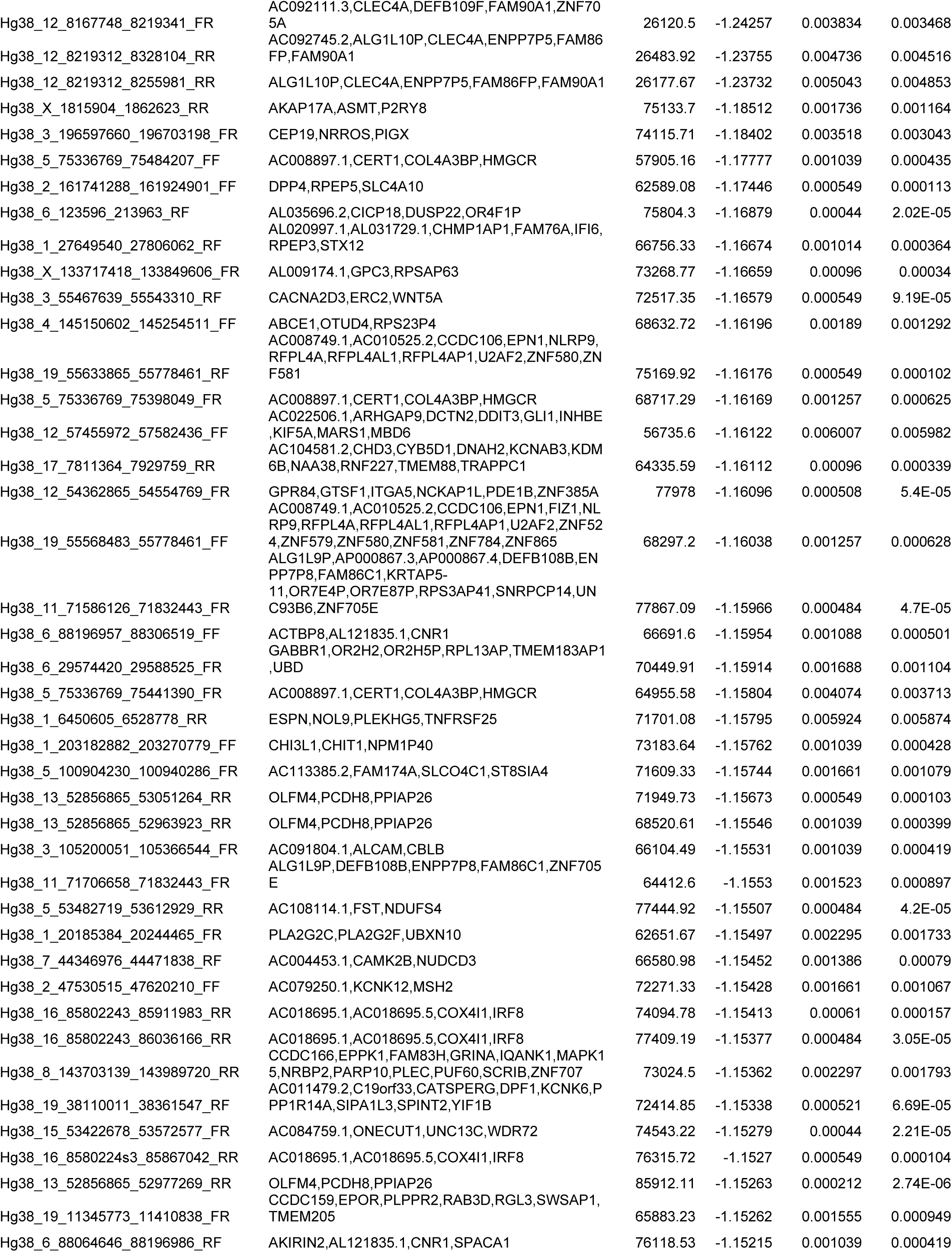
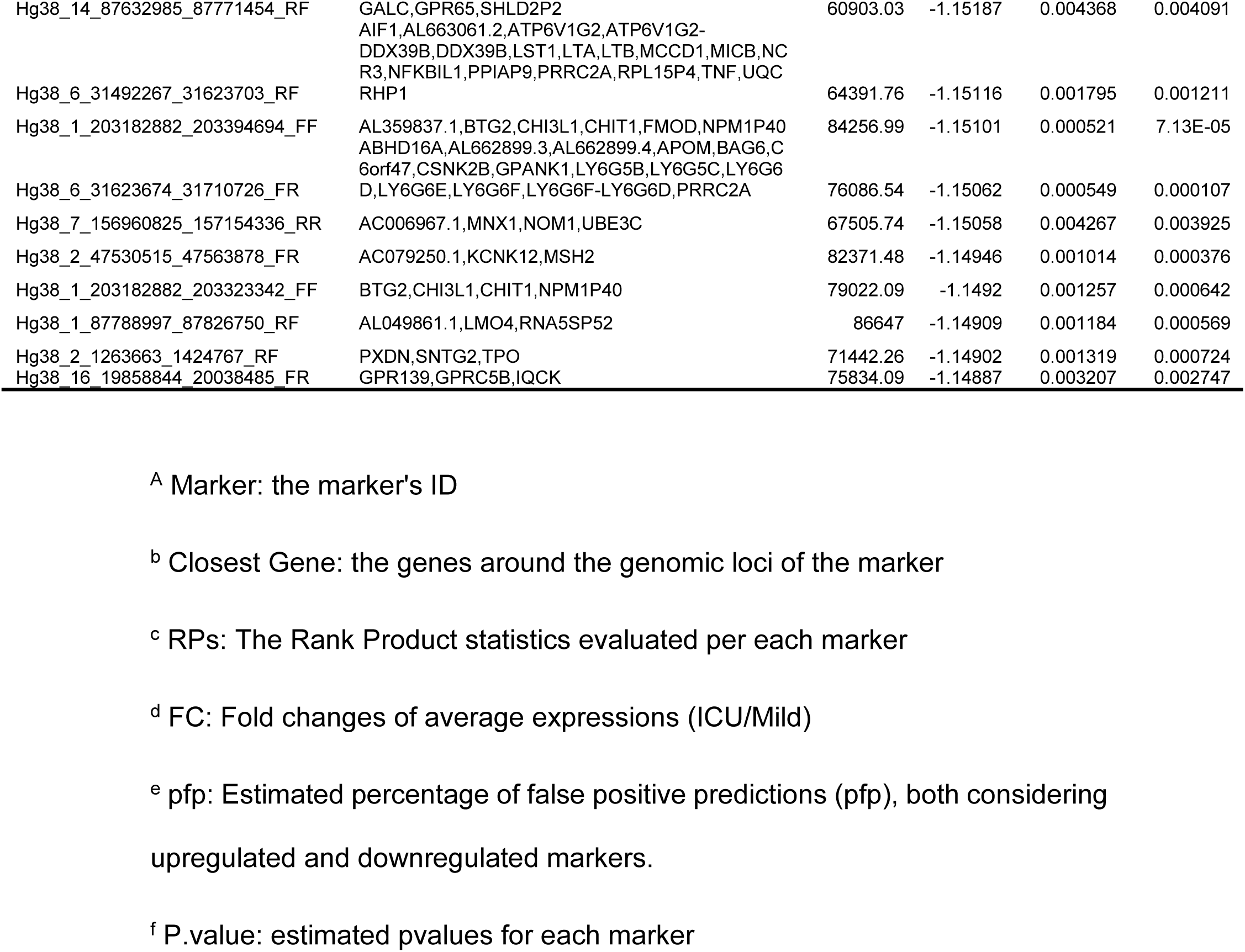
Top 100 3D genomic markers associated with Mild clinical outcomes.

#### Biological network analysis and therapeutic implications

Ranking top 100 immune-related 3D genomic markers associated with severe (ICU) outcome by adjusted p-value, then by abundance, we found the top 20 markers to be (**Table 2**) to be at genetic loci involved in macrophage-stimulating protein (MSP)-RON signalling (KLK5, NOS2, KLK3), G-Beta Gamma (Gβγ) Signalling (WNT2B, NOS2, VEGFC) and pathways related to regulation of nitric oxide. The top 20 3D genomic markers associated with Mild clinical outcomes in COVID-19 (**Table 2**) are at genomic loci involved in the regulation of RAC1 activation (PREX1, ARHGAP9), MHC II antigen presentation (KIF5A, DCTN2) and MHC 1 mediated antigen processing and presentation (FCGR1B, DCTN2, KIF5A). Interestingly, RAC1 signalling negatively regulates T cell migration via TCR signalling and inhibiting RAC1 restores T cell migration suggesting that essential mechanisms for T cell control are lacking in patients with Severe clinical presentations of COVID-19 [60].

When we mapped the genomic location of the top 100 3D genomic markers associated with Severe (ICU) clinical outcomes, we found broad genomic distribution with a notable high density at regions on chromosomes 5, 17, 20 and 22 (**Figure 4**).

**Figure 4.**
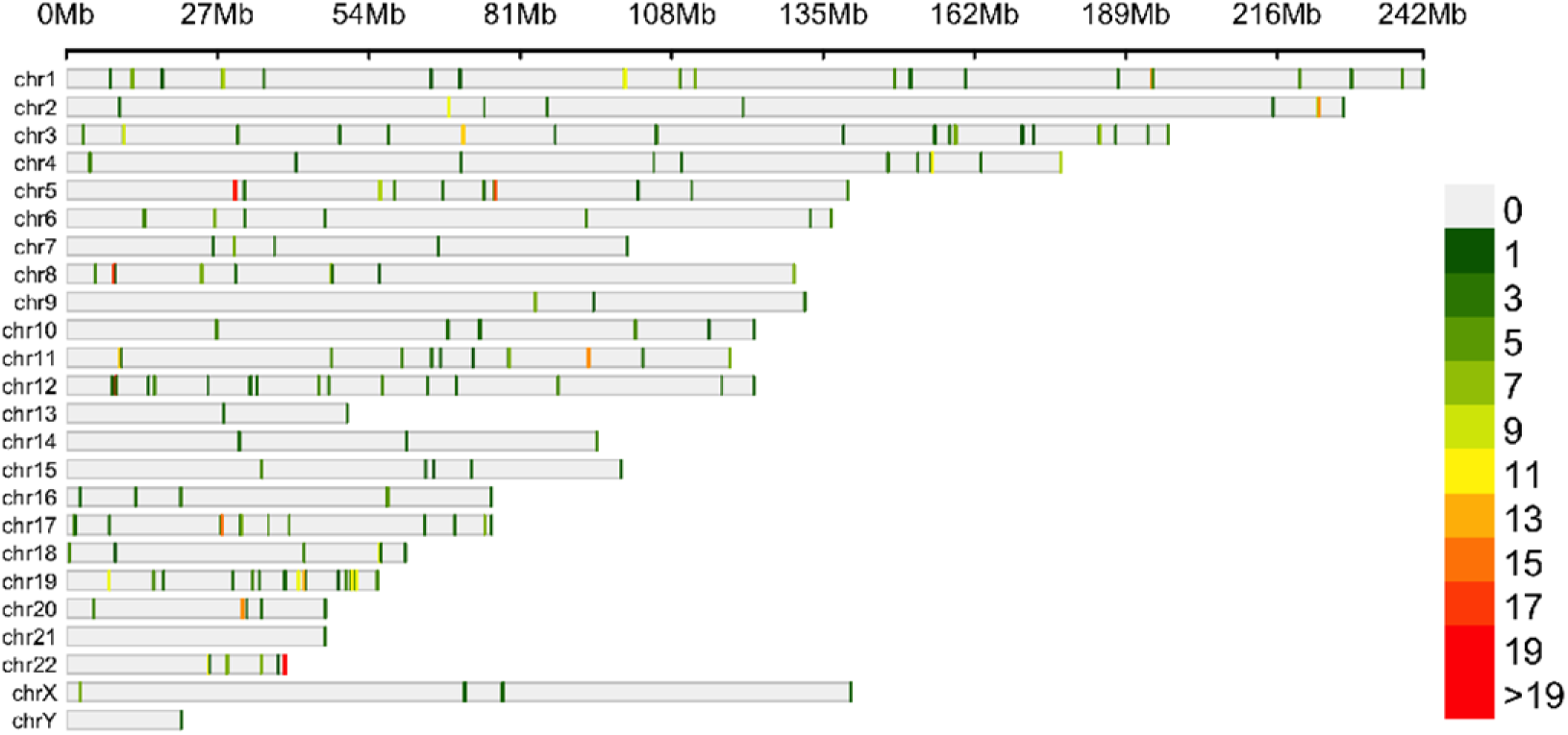
Genome wide mapping of 3D genomic loci associated with severe COVID-19 outcomes. Genomic locations and distribution of the top 100 3D genomic markers for Severe (ICU) clinical outcome. Individual human chromosomes are shown on the y-axis (chr1-chr22 along with the X and Y sex chromosomes). The heatmap shows the number of markers within a 0.3Mb genomic window with green representing a low density of markers and red indication a high density of markers. Sites with a high density of markers are seen on chromosomes 5, 17, 20 and 22.

Analysis of the top 3D genomic markers associated with Severe (ICU) COVID-19 outcomes using the Search Tool for Retrieval of Interacting Genes (STRING) database, we found a protein-protein interaction network with hubs on TNF, IL6, VEGFA, INS, TLR6, STAT1, MAPK1 and MAPK3 (**Supplementary Figure 1 and Supplementary Table 2)**.

Next, we used the network of genes under 3D genomic control for susceptibility to severe COVID-19 outcome to evaluate the existing drugs with known gene targets. We were trying to evaluate them as potential therapeutic tools for mitigation of severe disease outcomes. Using GeneAnalytics we identified 25 drug candidates with potential utility for treating COVID-19 disease (**Table 3**). Interestingly, the analysis based on 3D genomic profiling of severe COVID-19 patients identified at the second highest score Dexamethasone, which has been now reported as beneficial in reducing mortality among severe patients [61].

**Table 3:**
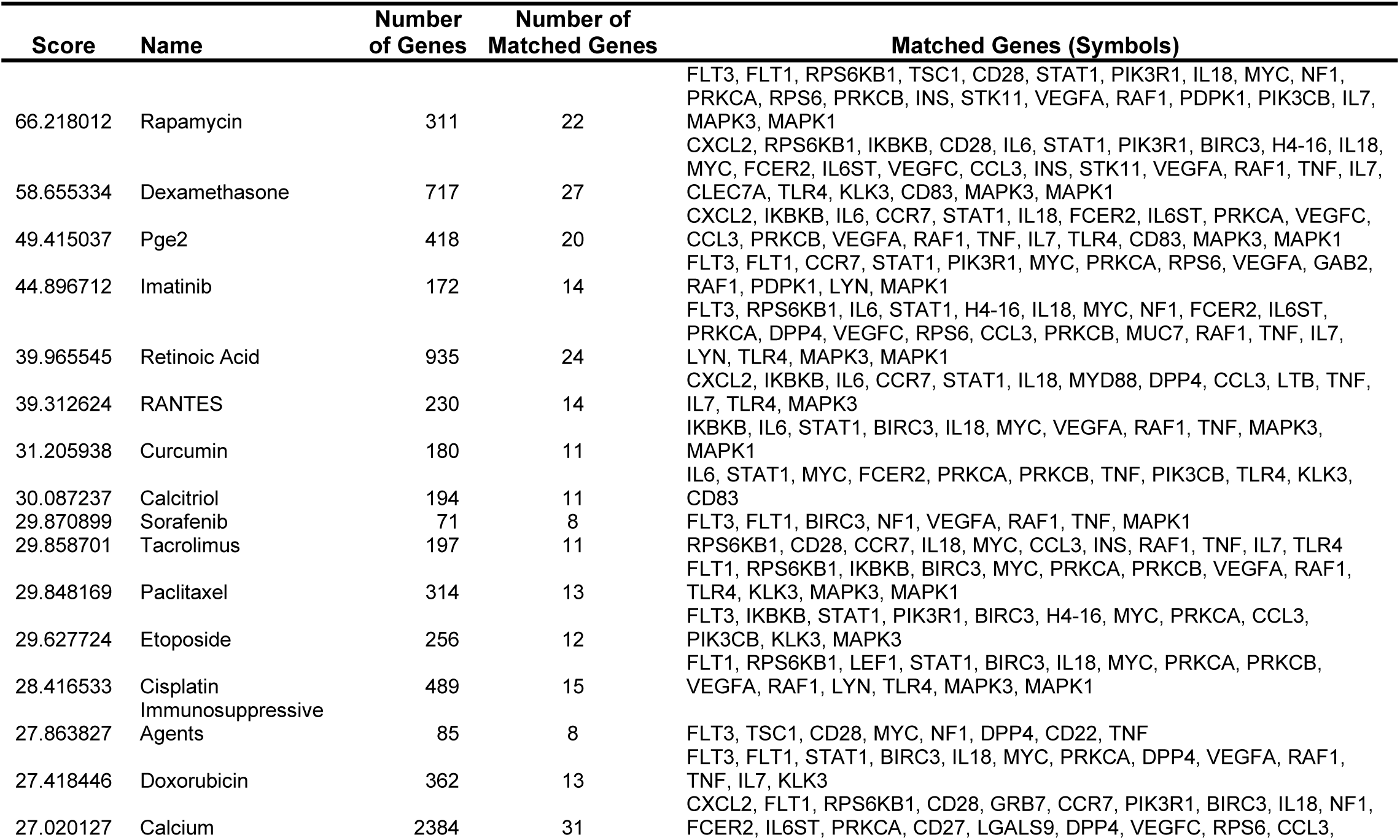
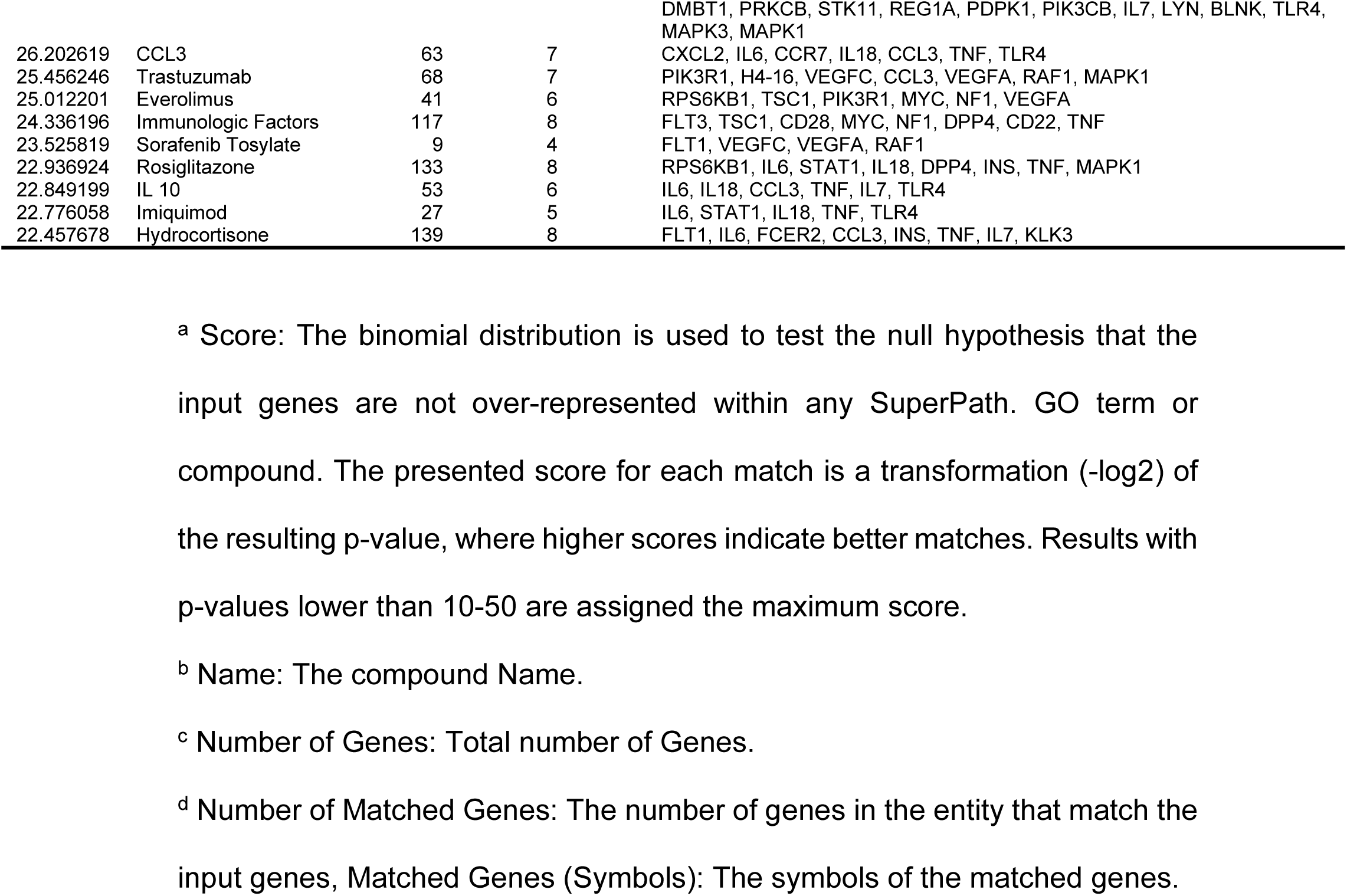
Potential Therapeutic Agent for Mitigation of Severe Disease Outcomes.

### Generation and testing of a blood-based 3D genomic predictive classifier for COVID-19 disease severity

Based on the established *EpiSwitch^®^* methodology for the discovery, evaluation, feature reduction and validation of 3D genome biomarkers in the format of a clinical assay [22–30], we wanted to evaluate the top 3D genomic marker leads identified here for further evidence of their prognostic stratification power before it could be developed into PCR-based *EpiSwitch^®^* test. The top 50 3D genomic markers associated with Severe (ICU) disease outcome were selected to build a classifier using machine learning algorithms at the level of *EpiSwitch^®^* array readouts. Markers associated with Severe (ICU) outcomes were selected over those associated with Mild clinical outcomes due to the tight clustering of the patient samples seen in the LDA analysis. Of the 80 patient samples used in this study, 38 samples (UK and USA) were peripheral blood mononuclear cells (PBMCs) and 42 (Peru) were whole blood collections. As *EpiSwitch^®^* clinical assays are developed on standard whole blood input [22–30], we used the 42 whole blood samples for classifier development and testing. Within these samples, 24 were Severe (ICU) patients and 18 were hospitalized patients with a Mild disease course. The 42 samples were randomly divided into a Training set of 30 samples and Test set of 12 samples. The use of both LDA and XGBoost algorithms to build classifier models was done to reduce the potential of data overfitting by either model. In addition to using multiple machine learning approaches to build classifying models, 10-fold cross validation was utilized to ensure robust and rigorous training models were built especially for XGBoost, this was carried out using the caret package in R. Both models were built using the top 50 3D genomic markers associated with Severe (ICU) outcomes. The LDA model built using the 30-patient Training set showed good separation by clinical outcome classes in both the Training and Test sets (**Figure 5A**).

**Figure 5.**
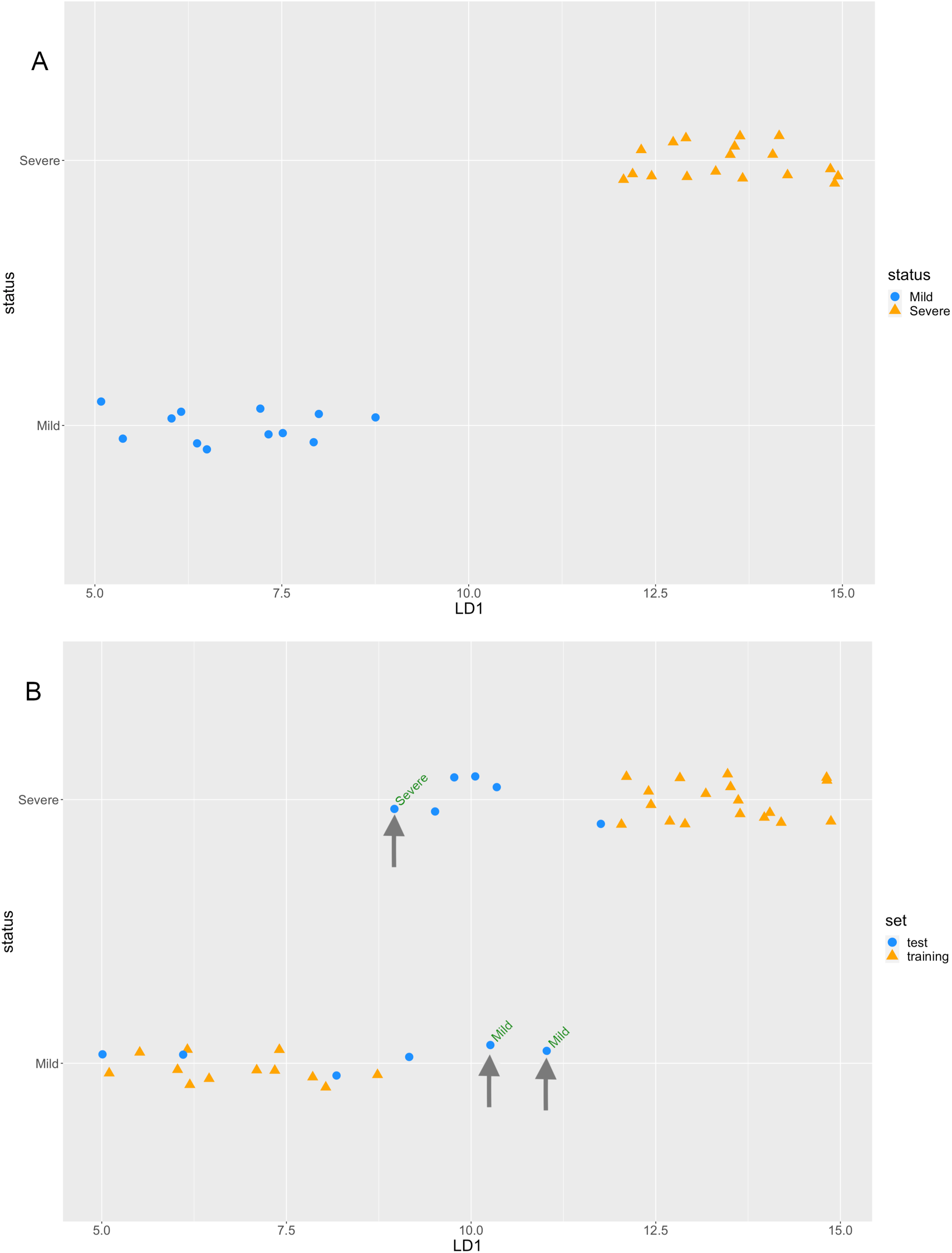
3D genomic markers can be used to classify COVID-19 disease outcomes. LDA plot of the Training (A) and Training and Test (B) sets of COVID-19 patients using the top 50 3D genomic markers associated with Severe (ICU) clinical outcomes. The markers showed separation of Mild (circles) and Severe (triangles) outcomes in Training Set (A), and across all 42 patients in training (triangle) and testing (circle) sets (B). Miscalled samples - 2 milds and 1 severe, are marked. Y axis - patient categories; X axis - Linear Discriminant Coordinate 1 (LD1).

As shown in Fig. 5, the LDA model miss-called 2 Mild patients as Severe and 1 Severe patient as Mild (**Figure 5B**). The overall confusion matrix for the overall Training and Test sample cohort of 42 patients is shown in **Figure 6**.

**Figure 6.**
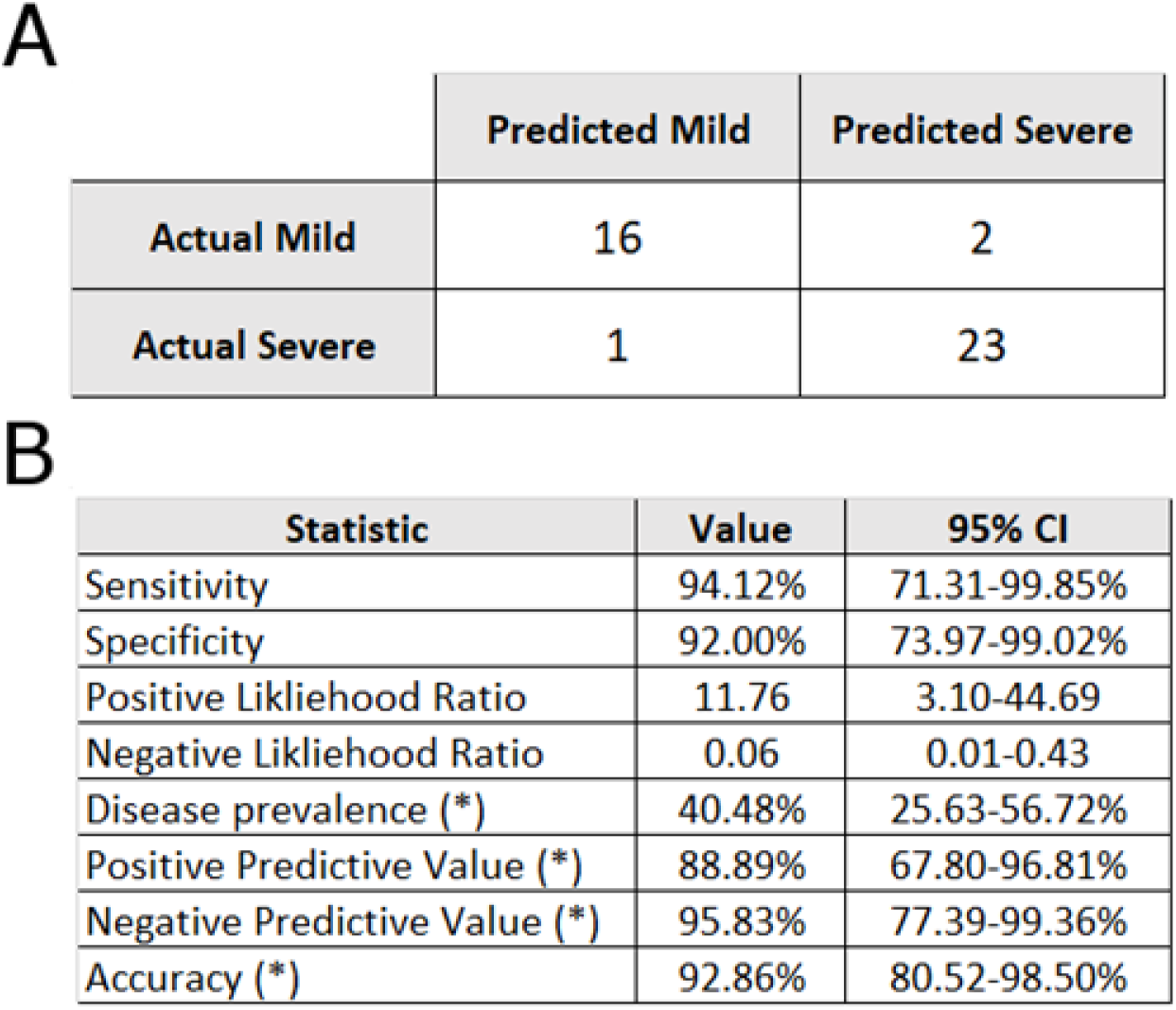
Confusion matrix and summary statistics for the LDA classifier. (A) LDA Classifier performance on the combined Training and Tests sets of 42 Mild and Severe (ICU) COVID-19 patients. (B) LDA classifier performance values for main test validation criteria and 95% confidence intervals (CI).

This LDA model result mirrors the classification of the Test samples using the model built on XGBoost (**Supplementary Figure 2**). Overall, both LDA and XGBoost models were able obtain a positive predictive value (PPV) of >83% for identifying ICU-severe patients with a balanced accuracy of >76%.

## Discussion

COVID infections lead to highly heterogeneous courses of COVID disease, from asymptomatic to mild to severe at considerable risk of fatality. With all of its genetic diversity and evolution, numerous mutations and genetic variants, including recent UK (B1.1.7) and South African (B.1.1.354) clades, the SARS-CoV-2 virus clearly shows strong variations in infectivity. However, there is little evidence for the virus to being responsible for heterogeneity of the disease [62]. Environmental factors, and largely the host itself, appears to be the major factor in defining disease severity and outcomes. As part of underlying mechanisms of heterogeneity, multiple studies point at the emerging evidence of differently pre-programed host innate immune cells in mild and severe disease courses, with strong differences in cellular responses occurring from the very early stages of infection [62].

Here, for the first time, we used an established 3D genomics approach to identify molecular biomarkers, known as chromosome conformation signatures (CCS), to capture the systemic differences in severe COVID-19 disease outcomes at the earliest stages of infection. The 3D genome acts as an integrator of multiple molecular inputs; from genetic risks, epigenetic modifications, transcriptional events and metabolic signalling and serves as a bridge between the stable conditional 3D genome and clinical outcomes [63, 64]. Preceding in the cascade of regulatory events the relevant and complex changes in gene expression, alterations in 3D genome structure represent a novel class of molecular readouts with the potential to forecast cellular and physiological phenotypes [63]. The *EpiSwitch* 3D genomic arrays used here have been reduced to a simple liquid biopsy testing platform with a demonstrated ability to provide diagnostic, prognostic, and predictive patient stratifications in a wide range of therapeutic areas [22–30], including COVID-19.

We used whole 3D genome arrays to analyse molecular profiles across 1.1 million sites per patient from 80 COVID-infected patients from around the world. In agreement with earlier observations, the regulatory 3D genome contained a stable network of 3D genomic alterations tightly associated with distinct clinical outcomes. The 3D genomic changes identified here provide further insight into functional networks of genome regulation after SARS-CoV-2 infection, as many were associated with genomic loci implicated in immune regulation. By biological network analysis, we identified several immune-related pathways that showed differential regulation in severe forms of COVID-19, including the (MSP)-RON pathway, Gβγ signalling, and nitric oxide (NO) signalling. MSP-RON signalling mediates systemic inflammatory responses and abnormalities in MSP-RON signalling can lead to onset of autoimmune disease [65]. Perhaps a more central mediator of the inflammatory response guiding the difference between mild and severe forms of COVID-19 identified by our study is Gβγ, which directly activates PI3Kγ and is involved in the recruitment of neutrophils to sites of inflammation. Inhibition of PI3Kγ significantly reduces inflammation and PI3Kγ is the common signalling effector for many different chemokine and receptors involved in promoting inflammation [66, 67]. Last, pro-inflammatory-macrophages upregulate inducible nitric oxide synthase (NOS2) and produce high steady state NO concentrations. NO is also responsible for modulating virtually all steps of innate and adaptive immunity [68, 69]. However, NO can also cause oxidative stress, which is especially damaging to the host due to increased tissue damage, a commonly seen pulmonary event seen in severe COVID-19 [2, 70, 71]. The cytokines IFN-γ and TNF-α, the latter of which was identified as a mediator of the severe clinical outcomes in COVID-19 in this study, are strong inducers of NOS2 [72]. Involvement of NO pathway may also explain higher risk in patients with microvascular pathologies: hypertension, diabetes, smoking, stroke, chronic kidney disease and age. It would be interesting to assess to what extent the respiratory failure in COVID patients is associated with hypoxia due to vascular mechanisms.

Our work has clinical diagnostic implications. Several cytokines and chemokines including TNF, MIP1α, and IL6 have been observed to be elevated using transcriptomics in patients with severe forms of COVID-19 [73, 74]. However, when measured at the protein level, these molecular factors aren’t elevated until after the onset of severe symptoms and are therefore unlikely to be causal in disease onset but may contribute to worsening of the severe disease. The 3D genomic changes on the other hand, represent one of the earliest steps in dynamic cellular phenotypic transitions and do not rely on longitudinal sampling during infection in order to predict disease severity outcome. We also identified some potentially novel therapeutic strategies for managing COVID-19. Interestingly, several of the drugs identified here as potential therapeutic tools have been tested independently in clinical trials for COVID-19, including mTOR inhibitors (rapamycin and tacrolimus) and general immunosuppressants (dexamethasone and hydrocortisone) [75–78]. In addition, we identified a potentially novel therapeutic pathway for COVID-19 that did not have evidence of being explored at the time of this writing based on a search of the ClinicalTrials.gov database. The signalling lipid prostaglandin E2 (PGE2), the cell signalling mediator calcium, the acute inflammatory phase cytokine CCL3 (also known as MIP1α) and the T-cell derived chemotactic cytokine CCL5 (also known as RANTES) were identified as potential pathways of therapeutic interest and intersect on a common signalling cascade. PGE2 exerts its cellular effects though binding to one of four cell membrane receptors (EP1-4) [79]. Binding to the EP1 or EP3 receptors increases intracellular calcium, while binding to EP2 and EP4 receptors triggers cyclic AMP mediated signalling events [79]. While PGE2 can act as a potent anti-inflammatory ligand, inhibiting the production of CCL3 in dendritic cells *in vivo* and the production of CCL5 mRNA and protein expression in LPS-activated macrophages *in vitro,* it can also be proinflammatory in certain lung conditions such as COPD, lung cancer, and several viral infections [80–82]. Elevated levels of PGE2 have been observed in SARS-CoV-2 infected patients and increased PGE2 has been postulated to correlate with enhanced COVID-19 severity in males [83, 84]. Although initial efforts at reducing PGE2 synthesis in COVID-19 through the use of non-steroidal anti-inflammatory drugs (NSAIDs) such as aspirin and ibuprofen have been controversial [85], our results suggest that prostaglandin signalling in immune cells may play an important role in mediating disease severity and should be evaluated as both a therapeutic intervention point as well as a consideration in clinical management of COVID-19 (e.g. patients with a history of NSAID use) [82].

Our findings also point to some logical next steps for advancing a clinically useful test for predicting COVID-19 severity. The 3D genomic markers identified by the bespoke *EpiSwitch^®^* Expoler array platform showed strong predictive power for COVID-19 disease outcome. Following previously published examples, these markers (or a subset of them) could be translated into a MIQE-compliant [22–30] qPCR-detectable format, reduced in number by machine learning on expanded patient cohorts, and used to develop a molecular classifier model with a minimal signature for validation on an independent testing cohort [22–30]. Interestingly, in our array study we found stratification of samples using either PBMC or whole blood collections highly consistent. The samples were collected at the time of COVID diagnosis, which took place from very early pre-symptomatic to advanced clinical manifestation stages of the disease. This suggests that the systemic 3D immune-genetic profiling, captured by *EpiSwitch^®^* technology, represents consistent features of regulatory state, including epigenetic status, prognostically associated with severe clinical outcome and represented in whole blood and PBMC samples.

With advance of vaccination programs around the world, the latest estimates from real-world broad immunization campaign in Israel demonstrate that for the age group >60 (89.9% received first dose and 80% received both doses of BNT162b2 BioNTech/Pfizer vaccine) hospitalization has dropped by 36% and severe cases by 29% [86, 87]. The latest Emergency Use Authorization of the single-shot Johnson &Johnson COVID-19 vaccine (JNJ-78436725) is based on the phase 3 ENSEMBLE trial results with overall 66% effectiveness in preventing moderate and severe disease, including 72% in US, 57% in Latin America, and 57% effectiveness in South Africa, tested against variant B.1.351 [88]. Together, this strongly suggests that in the immediate future, even in the context of active immunization, prognosis of COVID severity stratification would remain a valuable risk-mitigation tool for a significant part of the population.

The objective of the current study was to interrogate whether the clinical outcomes of severe versus mild profiles could be distinguished prognostically through the use of 3D genomics as a biomarker modality. As a secondary goal, we sought to use 3D genomics to better understand regulatory networks involved in the development of COVID-19 illness to aid in development or repurposed use of known therapeutic agents to control and mitigate severe outcomes. Open access to the data obtained here and direct access to *EpiSwitch^®^* Explorer array kits used in this study (**Materials and Methods**) should provide the research community with tools to further refine COVID-19 disease profiles and help better understand disease risk profiling towards the broader goal of promoting improved and informed clinical decision making.

## Conclusions

While the collective understanding of the biology behind SARS-CoV-2 infection leading to development of COVID-19 has been a focus of recent research efforts, the molecular factors that underly differences in COVID-19 disease presentation/severity are largely unknown and largely dictated by the host. There is a high need for high content technologies that could be reduced to practice for assessment of cellular, systemic host profiles to identify potential biomarkers and underlying molecular network mechanisms addressing outcome prediction. Taking an array-based *EpiSwitch^®^* approach, we identified blood-based differences in 3D immuno-genetic molecular profiles of patients with mild forms of COVID-19 compared to those with more severe clinical outcomes. Our results suggest that 3D immuno-genetic profiles in multi-cohort study could be used as a biomarker modality for COVID specific disease classification and outcome prediction.

In line with established *EpiSwitch^®^* methodology [22–30], further feature reduction in PCR format for the initial findings and biomarker leads described here, use of an extended patient cohort and an independent cohort validation will serve to further validate the biological relevance of the 3D genomic changes and could help to establish the minimal set of prognostic markers that could robustly stratify patients. Ultimately, as a next step, a reduced set of markers in *EpiSwitch^®^* MIQE-compliant qPCR format could be developed and used as a clinical prognostic assay to help patients understand their likely risk of developing severe disease outcomes if infected with the SARS-CoV-2 virus. Such a test could assist in informed clinical decisions on patient care in cases of a confirmed SARS-CoV-2 infection. It could also act as a prognostic measure, providing risk evaluation through systemic profiling prior to exposure to SARS-CoV-2 and assist in informed decisions on personal lifestyle, social, and workforce activities.

## Declarations

C.K., A.W., F.S., M.S., J.W. E.H. and A.A. are full-time employees at Oxford BioDynamics plc and have no other competing financial interests.

## Author Contributions

EH, AA conceived the study, MS, EH and AR assisted with *EpiSwitch^®^* array design. WM, AB and ZL assisted with design of experiments. MS and AA planned and reviewed experiments. CK, AW, FCS and EH analysed the data. RP, AD, PB, BE performed experiments. RV, AB, DP, PAR, JM helped with interpretation of the data and writing of the manuscript. WW, CK, EH, and AA. wrote and reviewed the manuscript.

## Acknowledgements

The authors would like to thank members of OBD Reference Facility Morgan Thacker, Louis James, Thomas Lavin, Catriona Williams, Matt Parnell, Aemilia Katzinski, Pieter Koorts, Kalsoom Rana and Jayne Green for direct operational support and taking part in processing of clinical samples, Olly Hunter for assistance in data analysis. In addition, we acknowledge Tissue Solutions, Boca Biolistics Inc., and BioIVT for the timely provision of high-quality clinical blood samples and clinical annotations, and Agilent Technologies, Inc. for supply of *EpiSwitch^®^* designed CGH microarray slides.

## Consent for publication

Written informed consent for publication was obtained from all authors.

## Availability of data and materials

The datasets used and/or analysed during the current study are available from the corresponding author on reasonable request.

## Ethics approval and consent to participate

All patients signed informed consent forms prior to providing blood samples. All ethical guidelines were followed.

---

**Supplemental Figure 1.**
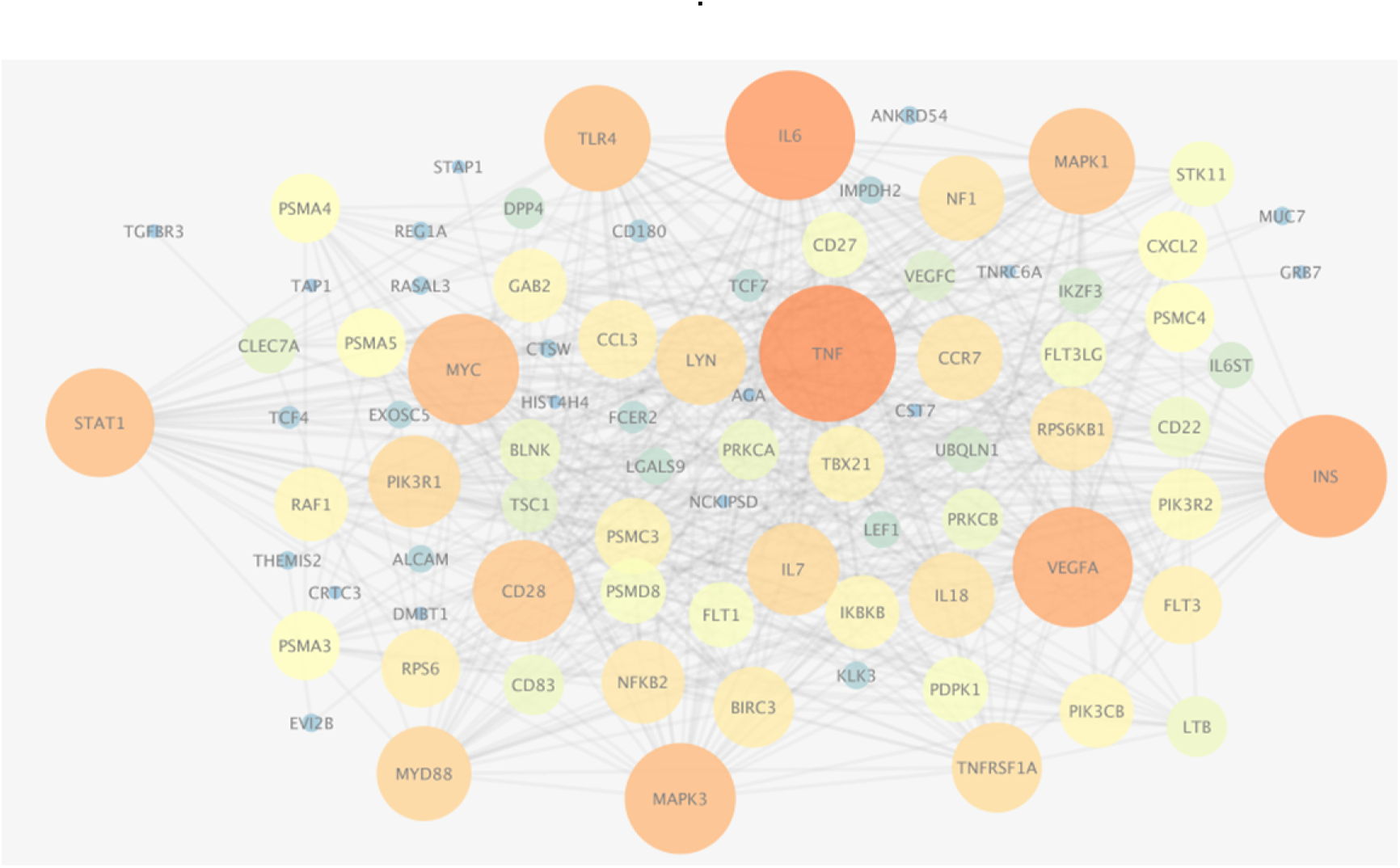
STRING Network associated with Severe (ICU) clinical outcomes in COVID-19. The proteins encoded by genes in the top 3D genomic markers associated with severe clinical outcomes in COVID-19 show a network with hubs at inflammatory mediators (TNF, IL6, VEGFA), immune-related receptors and signalling mediators (TLR4, STAT1, MAPK1) and the pleiotropic transcription factor MYC. See also Supplementary Table 2.

**Supplemental Figure 2.**
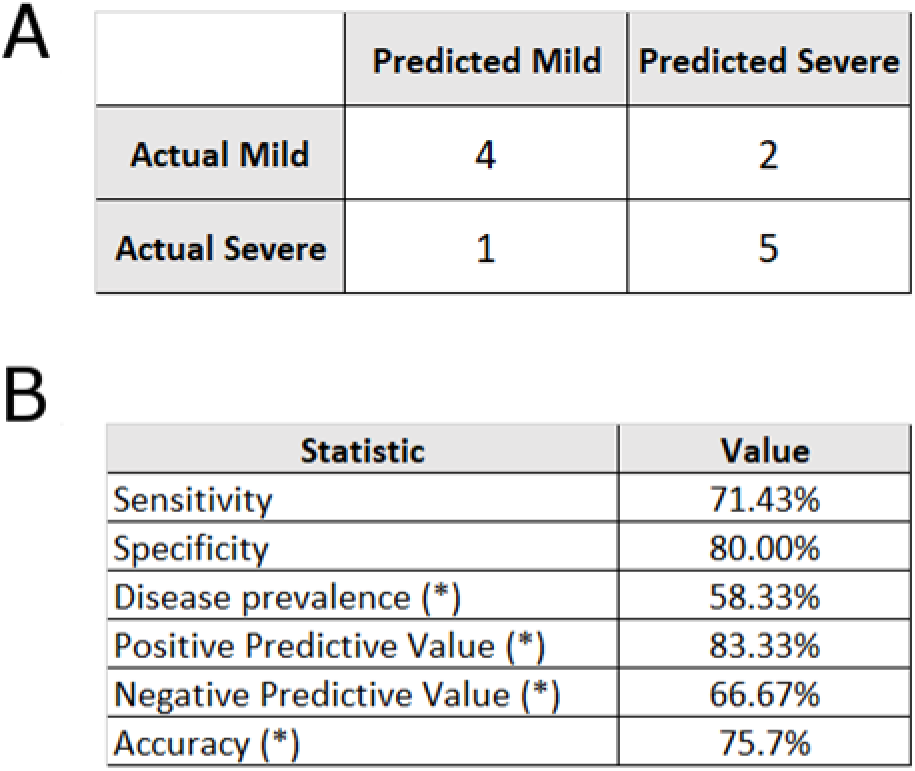
Confusion matrix and summary statistics for the XGBoost classifier. (A) XGBoost classifier performance on the Test set of 12 Mild and Severe (ICU) COVID-19 patients. (B) XGBoost classifier performance values for main test validation criteria and 95% confidence intervals (CI).

