## Supplementary figures and images for "3D genomic capture of regulatory immuno-genetic profiles in COVID-19 patients for prognosis of severe COVID disease outcome"

### Supplementary Figure 1

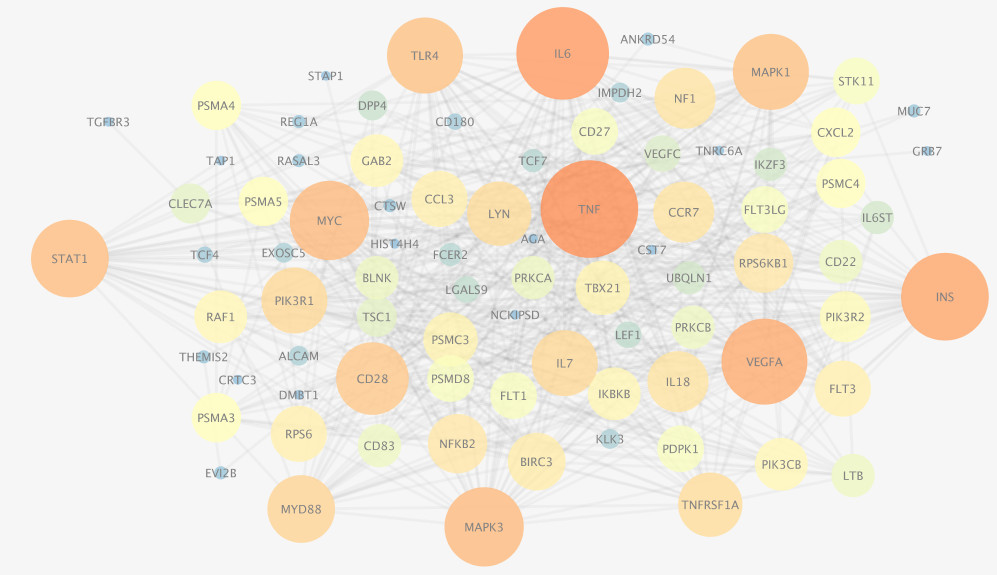

### Supplementary Figure 2

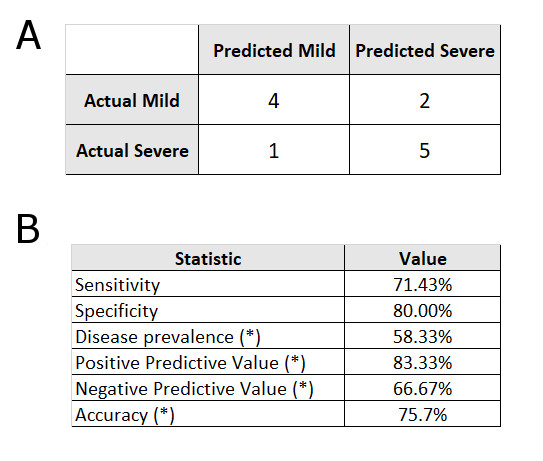
